# Relative Citation Ratio (RCR): A new metric that uses citation rates to measure influence at the article level

**DOI:** 10.1101/029629

**Authors:** B. Ian Hutchins, Xin Yuan, James M. Anderson, George M. Santangelo

## Abstract

Despite their recognized limitations, bibliometric assessments of scientific productivity have been widely adopted. We describe here an improved method that makes novel use of the co-citation network of each article to field-normalize the number of citations it has received. The resulting Relative Citation Ratio is article-level and field-independent, and provides an alternative to the invalid practice of using Journal Impact Factors to identify influential papers. To illustrate one application of our method, we analyzed 88,835 articles published between 2003 and 2010, and found that the National Institutes of Health awardees who authored those papers occupy relatively stable positions of influence across all disciplines. We demonstrate that the values generated by this method strongly correlate with the opinions of subject matter experts in biomedical research, and suggest that the same approach should be generally applicable to articles published in all areas of science. A beta version of iCite, our web tool for calculating Relative Citation Ratios of articles listed in PubMed, is available at https://icite.od.nih.gov.

## Introduction

In the current highly competitive pursuit of research positions and funding support (1), faculty hiring committees and grant review panels must make difficult predictions about the likelihood of future scientific success. Traditionally, these judgments have largely depended on recommendations by peers, informal interactions, and other subjective criteria. In recent years, decisionmakers have increasingly turned to numerical approaches such as counting first or corresponding author publications, using the impact factor of the journals in which those publications appear, and computing Hirsch or H-index values (2). The widespread adoption of these metrics, and the recognition that they are inadequate (3–6), highlight the ongoing need for alternative methods that can provide effectively normalized and reliable data-driven input to administrative decision-making, both as a means of sorting through large pools of qualified candidates, and as a way to help combat implicit bias.

Though each of the above methods of quantitation has strengths, accompanying weaknesses limit their utility. Counting first or corresponding author publications does on some level reflect the extent of a scientist’s contribution to their field, but it has the unavoidable effect of privileging quantity over quality, and may undervalue collaborative science (7). Journal impact factor (JIF) was for a time seen as a valuable indicator of scientific quality because it serves as a convenient, and not wholly inaccurate, proxy for expert opinion (8).

However, its blanket use also camouflages large differences in the influence of individual papers. This is because impact factor is calculated as the average number of times articles published over a two-year period in a given journal are cited; in reality, citations follow a log-normal rather than a Gaussian distribution (9). Moreover, since practitioners in disparate fields have differential access to high-profile publication venues, impact factor is of limited use in multidisciplinary science-of-science analyses. Despite these serious flaws, JIF continues to have a large effect on funding and hiring decisions (4,10,11). H-index, which attempts to assess the cumulative impact of the work done by an individual scientist, disadvantages early career stage investigators; it also undervalues some fields of research by failing to normalize raw citation counts (6).

Many alternative methods for quantifying scientific accomplishment have been proposed, including citation normalization to journals or journal categories (12–18), citation percentiles (14,19), eigenvector normalization (20,21), and source-normalization (13,22); the latter includes both the Mean Normalized Citation Score (17) and Source-Normalized Impact per Paper metrics (15,17,21–25). Although some of these methods have dramatically improved our theoretical understanding of citation dynamics (26–29), none have been widely adopted. To combine a further technical advance with a high likelihood of widespread adoption by varied stakeholders, including scientists, administrators and funding agencies, a new citation metric must overcome several practical challenges. From a technical standpoint, a new metric must be article-level, field-normalized in a way that is scalable from small to large portfolios without introducing significant bias at any level, and correlated with expert opinion. From an adoption standpoint, it should be freely accessible, calculated in a transparent fashion, and benchmarked to peer performance in a way that facilitates meaningful interpretation. Such an integrated benchmark, or comparison group, is not used by any currently available citation-based metric. Instead, all current measures aggregate articles from researchers across disparate geographical regions and institutional types, so that, for example, there is no easy way for primarily undergraduate institutions to directly compare the work they support against that of other teaching-focused institutions, or for developing nations to compare their research output to that of other developing nations (30). Enabling these and other apples-to-apples comparisons would greatly facilitate decision-making by research administrators.

We report here the development and validation of the Relative Citation Ratio (RCR) metric, which is based on the novel idea of using each article’s cocitation network to field-and time-normalize the number of citations it has received; this topically linked cohort is used to derive an expected citation rate, which serves as the ratio’s denominator. As is true of other bibliometrics, Article Citation Rate (ACR) is used as the numerator. Unlike other bibliometrics, though, RCR incorporates a customizable benchmarking feature that relates field-and time-normalized citations to the performance of a peer comparison group. RCR also meets or exceeds the standards set by other current metrics with respect to the ambitious ideals set out above. We use the RCR metric here to determine the extent to which National Institutes of Health (NIH) awardees maintain high or low levels of influence on their respective fields of research.

## Results

### Co-citation networks represent an article’s area of influence

Choosing to cite is the long-standing way in which one scholar acknowledges the relevance of another’s work. However, the utility of citations as a metric for quantifying influence has been limited, primarily because it is difficult to compare the value of one citation to another; different fields have different citation behaviors and are composed of widely varying numbers of potential citers (31,32). An effective citation-based evaluative tool must also take into account the length of time a paper has been available to potential citers, since a recently published article has had less time to accumulate citations than an older one. Finally, fair comparison is complicated by the fact that an author’s choice of which work to cite is not random; a widely known paper is more likely to be referenced than an obscure one of equal relevance. This is because the accrual of citations follows a power law or log-normal pattern, in accordance with a process called preferential attachment (26,32,33). Functionally this means that, each time a paper is cited, it is *a priori* more likely to be cited again.

An accurate citation-based measure of influence must address all of these issues, but we reasoned that the key to developing such a metric would be the careful identification of a comparison group, i.e., a cluster of interrelated papers against which the citation performance of an article of interest, or reference article (RA), could be evaluated. Using anetwork of papers linked to that Reference Article through citations occurred to us as a promisingpossibility (**Figure 1**). There are *a priori* three types of article-linked citation networks (34). A citing network is the collection of papers citing the Reference Article (**Figure 1a**, top row), a co-citation network is defined as the other papers appearing in the reference lists alongside the Reference Article (**Figure 1a**, middle row), and a cited network is the collection of papers in the reference list of the Reference Article (**Figure 1a**, bottom row).

**Figure 1.**
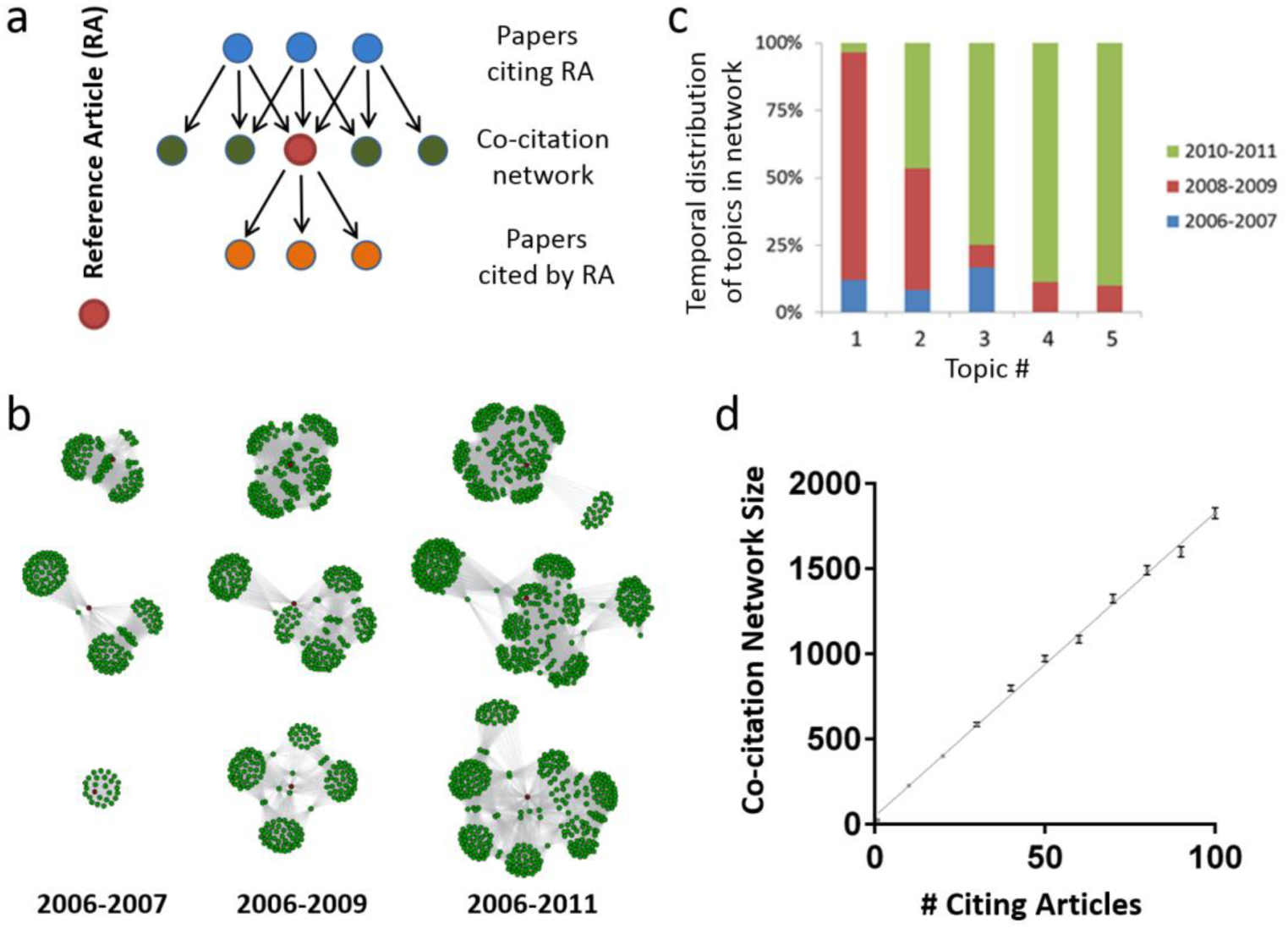
Properties of co-citation networks. (**a**) Schematic of a co-citation network. The Reference Article (RA) (red, middle row) cites previous papers from the literature (orange, bottom row);subsequent papers cite the RA (blue, top row). The co-citation network is the set of papers that appear alongside the article in the subsequent citing papers (green, middle row). The Field Citation Rate is calculated as the mean of the latter articles’ journal citation rates. (**b**) Growth of co-citation networks over time. Three RAs published in 2006 (red dots) were cited 5 (top row), 9 (middle row),or 31 times (bottom row) by 2011. Three intervals were chosen to illustrate the growth of the corresponding co-citation networks: 2006-2007, 2006-2009, and 2006-2011 (the first, second, and third columns, respectively). Each article in one of the three co-citation networks is shown as aseparate green dot; the edges (connections between dots) indicates their presence together in the same reference list. (**c**) Cluster algorithm-based content analysis of the 215 papers in the co-citation network of a sample reference article (RA; panel b, bottom network series) identified a changing pattern of relevance to different sub-disciplines over time. This RA described the identification of new peptides of possible clinical utility due to their similarity to known conotoxins. Papers in the co-citation network of this RA focused on: (1) α-conotoxin mechanisms of action; (2) structure and evolution of conotoxins; (3) cyclotide biochemistry; (4) conotoxin phylogenetics; and (5)identification and synthesis of lantibiotics. (**d**) Growth of an article’s co-citation network is proportional to the number of times it has been cited. Each point is the average network size of1000 randomly chosen papers with between 1 and 100 citations (error bars represent the standarderror of the mean). Each paper is only counted once, even if it is co-cited with the article of interest multiple times. An average of 17.8 new papers is added to the co-citation network for each additional citation. This suggests substantial duplication of articles within a co-citation network,since on average 32.4 papers (median of 30) are referenced in each citing article.

All three types of networks would be expected to accurately reflect the interdisciplinary nature of modern biomedical research and the expert opinion of publishing scientists, who are themselves the best judges of what constitutes a field. By leveraging this expertise, networks define empirical field boundaries that are simultaneously more flexible and more precise than those imposed by traditional bibliometric categories such as “biology and biochemistry” or “molecular biology”. An analysis of the co-citation network of a sample Reference Article illustrates this point. The Reference Article in the bottom panel of **Figure 1b** describes the identification of new peptides structurally similar to conotoxins, a little-known family of proteins that has begun to attract attention as the result of recent work describing their potential clinical utility (35). Although the papers in this network are all highly relevant to the study of conotoxins, they cross traditional disciplinary boundaries to include such diverse fields as evolutionary biology, structural biology, biochemistry, genetics, and pharmacology (**Figure 1c**).

Unlike cited networks, citing and co-citation networks can grow over time, allowing for the dynamic evaluation of an article’s influence; as illustrated by the example above, they can also indicate whether or not an article gains relevance to additional disciplines (**Figure 1b**, **c**). An important difference between citing and co-citation networks, however, is size. Since papers in the biomedical sciences have a median of 30 articles in their reference list, each citation event can be expected to add multiple papers to an article’s co-citation network (**Figure 1d**), but only one to its citing network. The latter are therefore highly vulnerable to finite number effects; in other words, for an article of interest with few citations, small changes in the citing network would have a disproportionate effect on how that article’s field was defined. We therefore chose to pursue co-citation networks as a way to describe an individual paper’s field.

Having chosen our comparison group, we looked for a way to test how accurately co-citation networks represent an article’s field. One way to characterize groups of documents is to cluster them based on the frequency at which specific terms appear in a particular document relative to the frequency at which they appear in the entire corpus, a method known as term-frequency-inverse document frequency (TF-IDF) (36). This is not a perfect approach, as it is possible to use entirely different words to describe similar concepts, but positive matches can be taken as a strong indication of similarity. Frequency of word occurrence can be converted into vectors, so that the cosine of the angle between two vectors is a measurement of how alike the two documents are. To evaluate co-citation networks relative to journal of publication, which is often used as a proxy for field, we selected more than 1300 articles with exactly five citations from six different journals and used cosine similarity analysis to compare their titles and abstracts to the titles and abstracts of each article in their co-citation network, and then separately to those of each article in the journal in which they appeared. Strikingly, this analysis showed that diagnostic words are much more likely to be shared between an article and the papers in its co-citation network than between that same article and the papers that appear alongside it in its journal of publication (**Figure 2**). As might be expected, the data in **Figure 2** also indicate that articles published in disciplinary journals are more alike than articles published in multidisciplinary journals.

**Figure 2.**
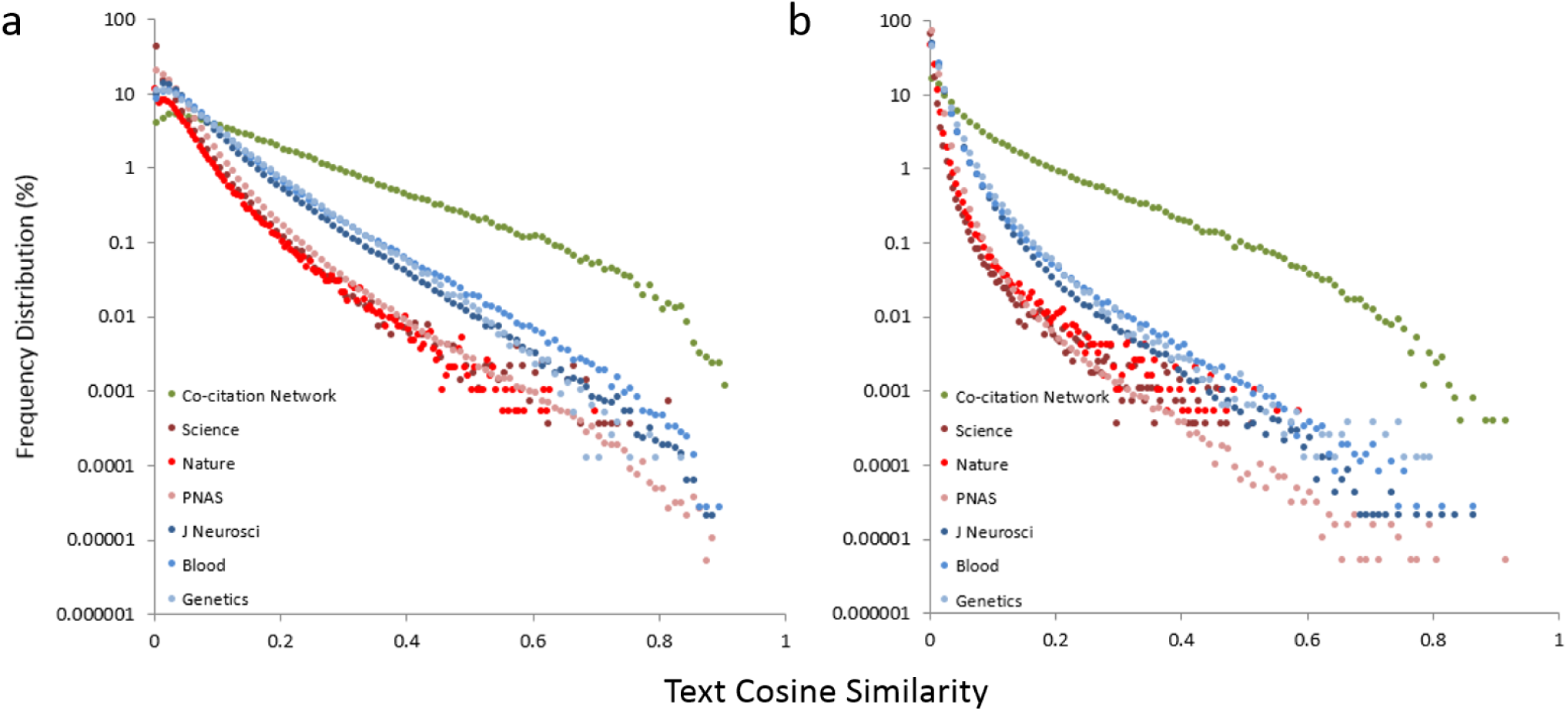
Text similarity of articles is defined more accurately by their co-citation networks than by the journals in which they appear. (**a**) The text in each of 1397 reference articles was compared, either with the text in each corresponding co-citation network, or separately with the collection of articles appearing in the same journal. Cosine similarity scores were then calculated, using either the top 1000 terms (**a**) or all terms appearing in at least 10 documents (**b**). Filled circles in green, co-citation network comparison; filled circles in shades of red, disciplinary journal comparison; filled circles in shades of blue, multidisciplinary journal comparison. Curves shifted to the right show more text similarity: reference articles are least similar to papers in the same multidisciplinary journals, more similar to papers in the same disciplinary journal, and most similar to papers in their co-citation network.

### Calculating the Relative Citation Ratio

After demonstrating that co-citation networks accurately represent an article’s field, our next step was to decide how to calculate the values that numerically represent the co-citation network of each Reference Article. The most obvious choice, averaging the citation rates of articles in the co-citation network, would also be highly vulnerable to finite number effects. We therefore chose to average the citation rates of the journals represented by the collection of articles in each co-citation network. If a journal was represented twice, its journal citation rate (JCR) was added twice when calculating the average JCR. For reasons of algorithmic parsimony we used the JCRs for the year each article in the co-citation network was published; a different choice at this step would be expected to have little if any effect, since almost all JCRs are quite stable over time (**Supporting Figure S1**; **Supporting Table S1**). Since a co-citation network can be reasonably thought to correspond with a Reference Article’s area of science, the average of all JCRs in a given network can be redefined as that Reference Article’s Field Citation Rate (FCR).

Using this method (**Figure 3**; **Supporting Figure S2**; **Supporting Equations S1 and S2**), we calculated Field Citation Rates for 35,837 papers published in 2009 by NIH grant recipients, specifically those who received R01 awards, the standard mechanism used by NIH to fund investigator-initiated research. We also calculated what the Field Citation Rate would be if it were instead based on citing or cited networks. It is generally accepted that, whereas practitioners in the same field exhibit at least some variation in citation behavior, much broader variation exists among authors in different fields. The more closely a method of field definition approaches maximal separation of between-field and within-field citation behaviors, the lower its expected variance in citations per year (CPY). Field Citation Rates based on co-citation networks exhibited lower variance than those based on cited or citing networks (Table 1), suggesting that co-citation networks are better at defining an article’s field than citing or cited networks. As expected, Field Citation Rates also display less variance than either Article Citation Rates (*p* < 10^−4^, F-test for unequal variance) or JIFs (*p* < 10^−4^, F-test for unequal variance, **Figure 3b**, **Table 1**).

**Figure 3.**
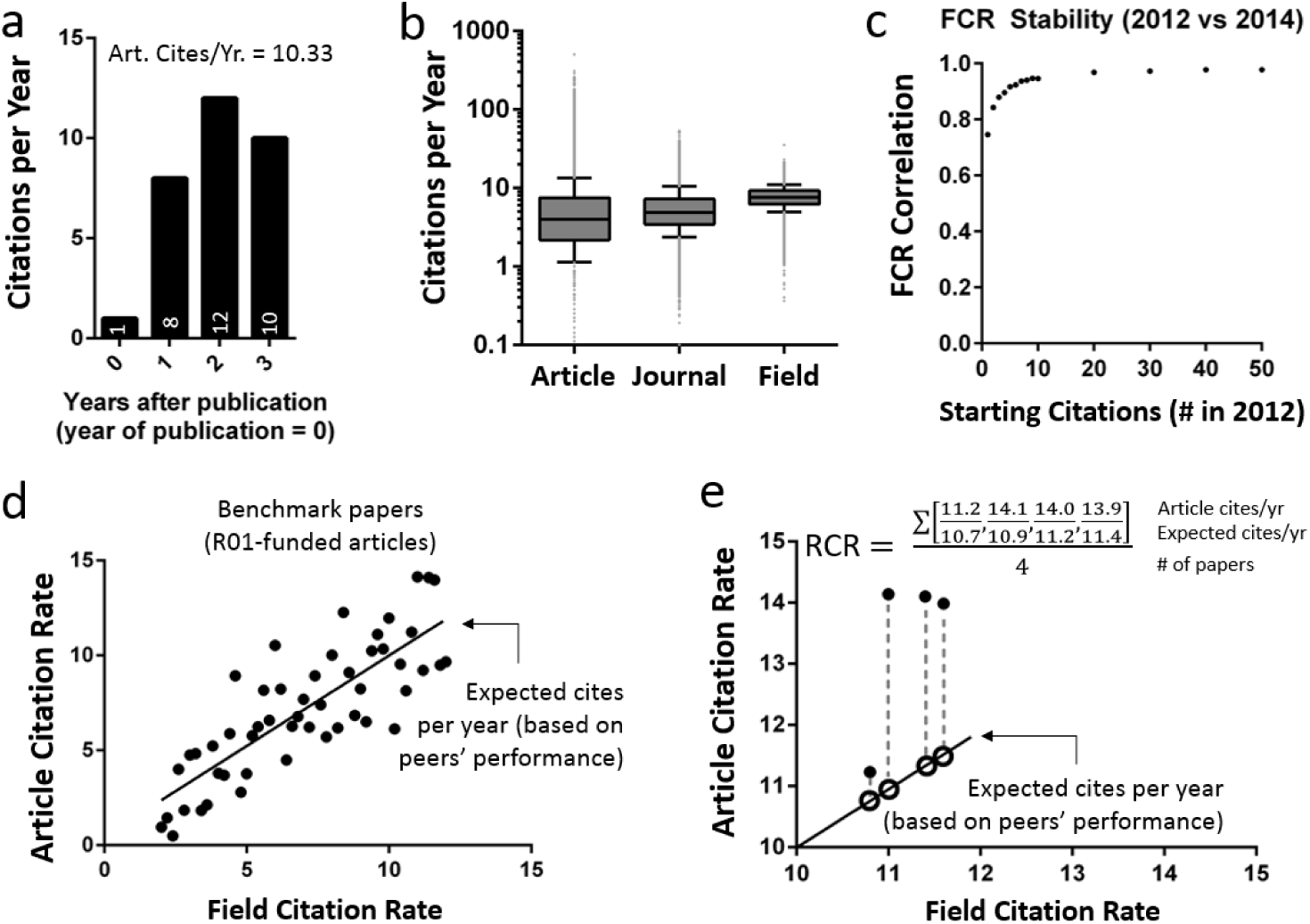
Algorithm for calculating the Relative Citation Ratio. (**a**) Article Citation Rate (ACR) is calculated as the total citations divided by the number of years excluding the calendar year of publication (**Supporting Equation S1**), when few, if any, citations accrue (**Supporting Figure S2**). (**b**) Box-and whisker plots of 88,835 NIH-funded papers (published between 2003 and 2010), summarizing their Article Citation Rate, Journal Impact Factor (matched to the article’s year of publication), and Field Citation Rate. Boxes show the 25th-75th percentiles with a line at the median; whiskers extend to the 10th and 90th percentiles. (**c**) Correlation of FCR as generated in 2012 vs. two years later in 2014 for the same set of articles, as a function of the number of starting citations in 2012. Data were sliced by the number of initial citations in 2012, to assess stability as a function of the number of citing articles (and thereby the starting size of the network). Each point, correlation coefficient for > 1000 articles. Between 2012 and 2014, articles accrued a median of 5 additional citations. (**d**) Generate an expectation for article citation rates based on a preselected benchmark group, by regressing the ACR of the benchmark papers onto their FCRs (**Supporting Equations S3, S4**), one regression each publication year. The graphed examples were sampled from a random distribution for illustrative purposes. (**e**) The coefficients from each year’s regression equation transforms the Field Citation Rates of papers published in the same year into Expected Citation Rates (**Supporting Equation S5**). Each paper’s RCR is its ACR/ECR ratio. A portfolio’s RCR is simply the average of the individual articles’ RCRs (**Supporting Equation S6**).

**Table 1.** Variance of Field Citation Rates and Expected Citation Rates using different levels of the citation network for calculations (based on 35,837 R01-funded papers published in 2009).

| Network level | Variance (FCR) | Variance (ECR) |
| --- | --- | --- |
| Impact Factor (RA only) | 33.7 | 30.8 |
| Cited network | 16.1 | 9.5 |
| Citing network | 6.7 | 7.8 |
| Co-citation network | 3.4 | 3.4 |

We next asked how stable the Field Citation Rates in our dataset remain over time, particularly where the starting co-citation network is small. To answer this question, we calculated Field Citation Rates for the 262,988 papers published by R01 grantees between 2003 and 2011 and cited one or more times through the end of 2012, then recalculated Field Citation Rates for the same articles, this time using citations accrued through the end of 2014 (**Figure 3c**). Comparison of the two values shows that earlier Field Citation Rates are well aligned with later ones, even when the initial cocitation network was built on a single citation (Pearson correlation coefficient *r* of 0.75 vs. two years later). The Field Citation Rate quickly converged within 5 citations, passing ***r*** = 0.9 at that point (**Figure 3c**). The consistency that Field Citation Rate values display should not be surprising, given the manner in which additional citations rapidly grow an article’s co-citation network (**Figure 1d**), each node of which represents a citation rate that is itself derived from a large collection of articles (**Supporting Equations S1 and S2**). In this way our method of calculation provides a low-variance quantitative comparator, while still allowing the articles themselves to cover a highly dynamic range of subjects.

Having established the co-citation network as a means of determining a Field Citation Rate for each Reference Article, our next step was to calculate ACR/FCR ratios. Since both Article Citation Rates and Field Citation Rates are measured in citations per year, this generates a rateless, timeless metric that can be used to assess the relative influence of any two Reference Articles. However, it does not measure these values against any broader context. For example, if two Reference Articles have ACR/FCR ratios of 0.7 and 2.1, this represents a three-fold difference in influence, but it is unclear which of those values would be closer to the overall mean or median for a large collection of papers. One additional step is therefore needed to adjust the raw ACR/FCR ratios so that, for any given Field Citation Rate, the average RCR equals 1.0. Any selected cohort of Reference Articles can be used as a standard for anchoring expectations, i.e. as a customized benchmark (**Supporting Equations S3–S6**). We selected R01-funded papers as our benchmark set; for any given year, regression of the Article Citation Rate and Field Citation Rate values of R01-funded papers yields the equation describing, for the Field Citation Rate of a given Reference Article published in that year, the expected citation rate (**Figure 3d** and **Supporting Table S2**). Inserting the Article Citation Rate as the numerator and Field Citation Rate of that Reference Article into the regression equation as the denominator is the final step in calculating its RCR value, which incorporates the normalization both to its field of research, and to the citation performance of its peers (**Figure 3e** and **Supporting Information**).

We considered two possible ways to regress Article Citation Rate on Field Citation Rate in the benchmarking step of the RCR calculation. The ordinary least squares (OLS) approach will benchmark articles such that the mean RCR is equal to 1.0. OLS regression is suitable for large-scale analyses such as those conducted by universities or funding agencies. However, in smaller analyses where the distribution of data may be skewed, OLS may yield an RCR less than 1.0 for the median, i.e. typical, article under consideration. In situations such as these, for example in the case of web tools enabling search and exploration at the article or investigator level, quantile regression is more desirable as it yields a median RCR equal to 1.0.

### Expert validation of RCR as a measure of influence

For the work presented here, we chose as a benchmark the full set of 311,497 Reference Articles published from 2002 through 2012 by NIH-R01 awardees. To measure the degree of correspondence between our method and expert opinion, we compared RCRs generated by OLS benchmarking of ACR/FCR values with three independent sets of postpublication evaluations by subject matter experts (details in **Supporting Information**). We compared RCR with expert rankings for 2193 articles published in 2009 and evaluated by Faculty of 1000 members (**Figure 4a** and **Supporting Figure S3**), as well as rankings of 430 Howard Hughes Medical Institute-or NIH-funded articles published between 2005 and 2011 and evaluated in a study conducted by the Science and Technology Policy Institute (STPI, **Figure 4b** and Supporting Figure S4), and finally, 290 articles published in 2009 by extramurally funded NIH investigators and evaluated by NIH intramural investigators in a study of our own design (**Figure 4c**; **Supporting Figures S5–S7**). All three approaches demonstrate that RCR values are well correlated with reviewers’ judgments. We asked experts in the latter study to provide, in addition to an overall score, scores for several independent sub-criteria: likely impact of the research, importance of the question being addressed, robustness of the study, appropriateness of the methods, and human health relevance. Random Forest analysis (37) indicated that their scores for likely impact were weighted most heavily in determining their overall evaluation (**Supporting Figure S6**).

**Figure 4.**
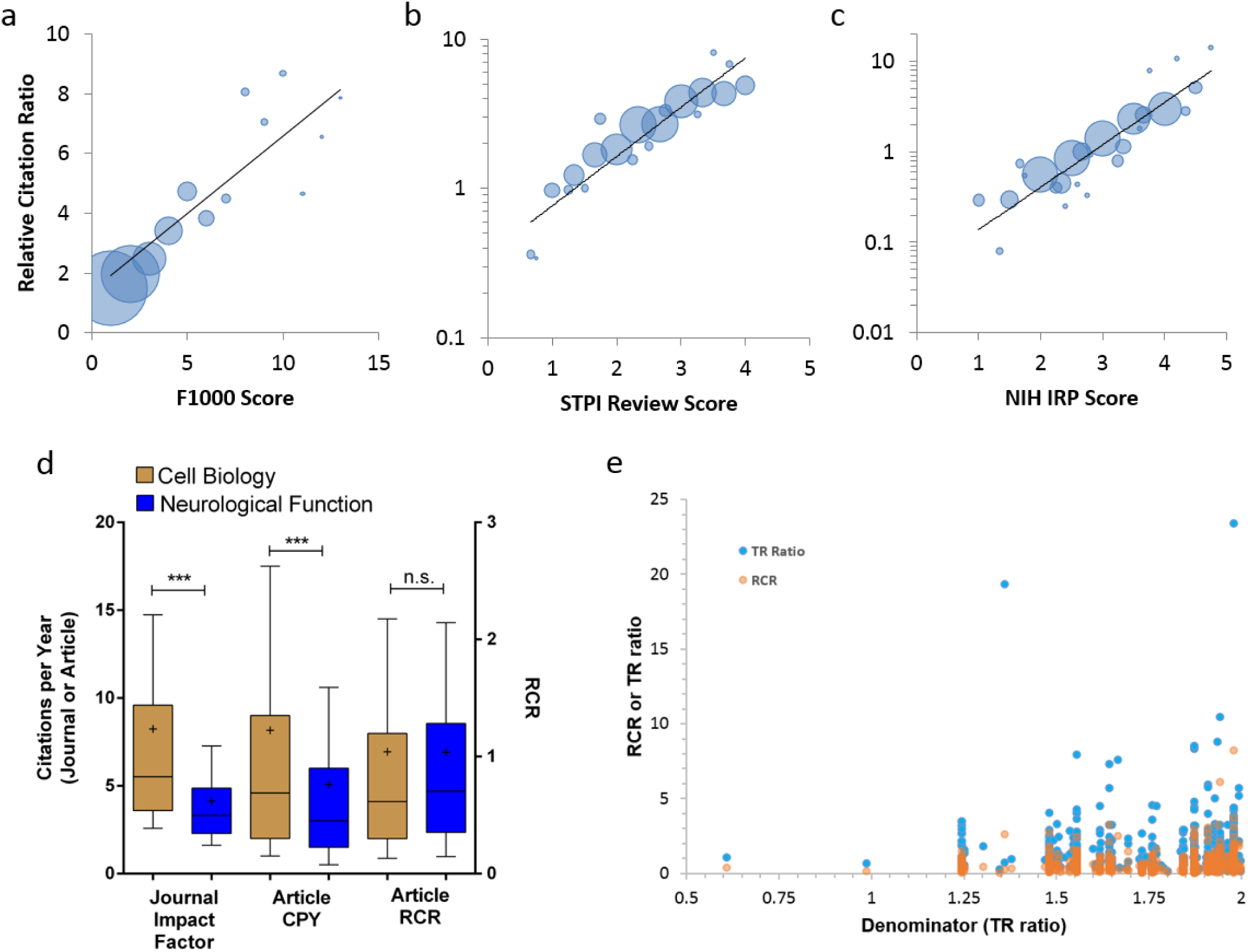
Relative Citation Ratios correspond with expert reviewer scores. (**a-c**) Bubble plots of reviewer scores vs. RCR for three different datasets. Articles are binned by reviewer score; bubble area is proportionate to the number of articles in that bin. (**a**) F1000 scores for 2193 R01-funded papers published in 2009. Faculty reviewers rated the articles on a scale of one to three (“Good”, “Very Good”, and “Exceptional”, respectively); those scores were summed into a composite F1000 score for each article (**Supporting Figure 3**). (**b**) Reviewer scores of 430 HHMI and NIH-funded papers collected by the Science and Technology Policy Institute. (**c**) Scores of 290 R01-funded articles reviewed by experts from the NIH Intramural Research Program. Black line, linear regression. (**d**) Box-and-whisker plots illustrating the distribution of journal impact factors (JIFs) citations per year (CPY) and relative citation ratios (RCR) for two areas of NIH funded research. Cell Biology, n = 5936; Neurological Function, n = 5417. *** p < 0.001, Kruskal-Wallis with Dunn’s multiple comparison test. n.s., not significant. Mean represented by a “*”. (**e**) Comparison of RCRs (orange) and Thompson Reuters ratios (blue; 17,27) for the same 544 articles with a low denominator. Data points are partially transparent to allow coordinates with multiple papers (darker) to be more clearly identified.

In addition to correlating with expert opinion, RCR is ranking invariant, which is considered to be a desirable property of bibliometric indicators (38,39). In short, an indicator is ranking invariant when it is used to place two groups of articles in hierarchical order, and the relative positions in that order do not change when uncited articles are added to each group. The RCR metric is ranking invariant when the same number of uncited articles is added to two groups of equal size (**Supporting Equations S7–S9**). RCR is also ranking invariant when the same proportion of uncited articles is added to two groups of unequal size (**Supporting Equations S10–S11**). This demonstrates that the RCR method can be used effectively and safely in evaluating the relative influence of large groups of publications.

### Comparison of RCR to existing bibliometrics

The ideal bibliometric method would provide decision makers with the means to perfectly evaluate the relative influence of even widely divergent areas of science. One way to test whether RCR represents a step towards this ambitious goal is to compare its performance in various scenarios to that of existing metrics. To begin, we asked whether RCR can fairly value a field that is disadvantaged by the use of two of the most widely recognized markers of influence: journal impact factor and citations per year. Two areas of science in which the National Institutes of Health funds research are neurological function and basic cell biology. Both subjects are deserving of attention and resources; however, papers in the former field, which includes subjects such as dementia and mental health, tend to appear in lower impact factor journals and receive fewer citations per year than those in the latter. In contrast, the distribution of RCR values for these two areas of study is statistically indistinguishable (**Figure 4d**). Although this is a single example, it does illustrate one way in which RCR provides value beyond either of these two alternative metrics.

While impact factor and citations per year are two of the most commonly used evaluative measures, they are also arguably less sophisticated than other field-normalized methods advanced by bibliometricians. A recent publication has reported that RCR is better correlated with expert opinion than one of these, MNCS, but slightly less well than another, SNCS_2_ (40). However, SNCS_2_ has an important disadvantage relative to RCR; it grows continually over time (like raw citation counts), and so is biased in favor of older papers. This disadvantage can be countered by calculating SNCS2 over a fixed time window, but doing so can obscure important dynamic aspects of citation behavior. In other words, if an article becomes more or less influential compared to its peers after that fixed window has passed, as for example can occur with “sleeping beauties” (41), this would not be apparent to users of SNCS_2_. These authors (40) also found that RCR is better correlated with expert opinion than citation percentiles, which is the measure bibliometricians have previously recommended as best for evaluating the impact of published work (42). A different team of researchers has also recently reported that a simplified version of the RCR algorithm is better at identifying important papers than Google’s PageRank (43), which has previously been adapted to quantitate the influence of an article or author (44–46).

One concern that arises whenever citations are field-normalized is that papers in disciplines with intrinsically low citation rates might be inappropriately advantaged. To evaluate how RCR meets this challenge, we compared our approach to an adaptation of the MNCS method, the Thomson-Reuters (TR) ratio. Like RCR, the TR ratio uses citations per year as its numerator; unlike RCR, the TR denominator is based on the average citation count of all articles published in a given year in journals that are defined as part of the same category (see **Supporting Information**). This journal categorization approach has some problems; bibliometricians have expressed concern that it is not refined enough to be compatible with article-level metrics (47), and in the case of TR ratios, the use of proprietary journal category lists renders the calculation of the metric somewhat opaque. Still, it is a more refined measure than JIF or CPY, and like RCR, it seeks to field-normalize citations based on choice of comparison group in its denominator.

A TR ratio is available for 34,520 of the 35,837 PubMed Indexed papers published in 2009 by recipients of NIH R01 grants. For this set of articles, the TR ratio denominator ranges from 0.61 to 9.0, and a value of 2.0 or less captures 544 papers (the bottom 1.6%; **Figure 4e**). The average TR ratio for these papers is 1.67, and the average RCR is 0.67. Since RCR is benchmarked so that the mean paper receives a score of 1.0, it is immediately obvious that these works are having relatively little influence on their respective fields, in spite of their intrinsically low field citation rates. In contrast, TR ratios are not benchmarked, so it is difficult to know whether or not it would be appropriate to flag these works as relatively low influence. Nevertheless, we can compare how each method ranks papers with these very low denominators by calculating the fraction with values above the RCR and TR ratio medians. Of the 544 low-denominator articles, 290 have a TR ratio greater than 1.07, the median value of the 34,520 papers, whereas 205 articles are above the RCR median of 0.63. This pattern holds when comparing the number of low-denominator articles that each approach places in the top 5% of the overall distribution; the TR method identifies seventeen such papers, whereas the RCR method identifies eight. Therefore, in avoiding inappropriate inflation of the assessment of articles with the lowest field citation rates, RCR is at least as good as, and arguably better than, MNCS.

Importantly, RCR is an improvement over existing metrics in terms of accessibility. While citation percentiles and TR ratios are only available through an expensive institutional subscription service, RCR values for PubMed indexed articles are freely available through the web-based *iCite* calculator, a screenshot of which is shown in **Figure 5**. For each PMID entered into *iCite*, users can download an Excel spreadsheet showing the total number of citations and the number of citations per year received by that publication; the number of expected citations per year, which are derived from a benchmark group consisting of all NIH R01 grantees, and the Field Citation Rate are also reported for each article. Detailed, step-by-step help files are posted on the *iCite* website, and the full code is available on GitHub.

**Figure 5.**
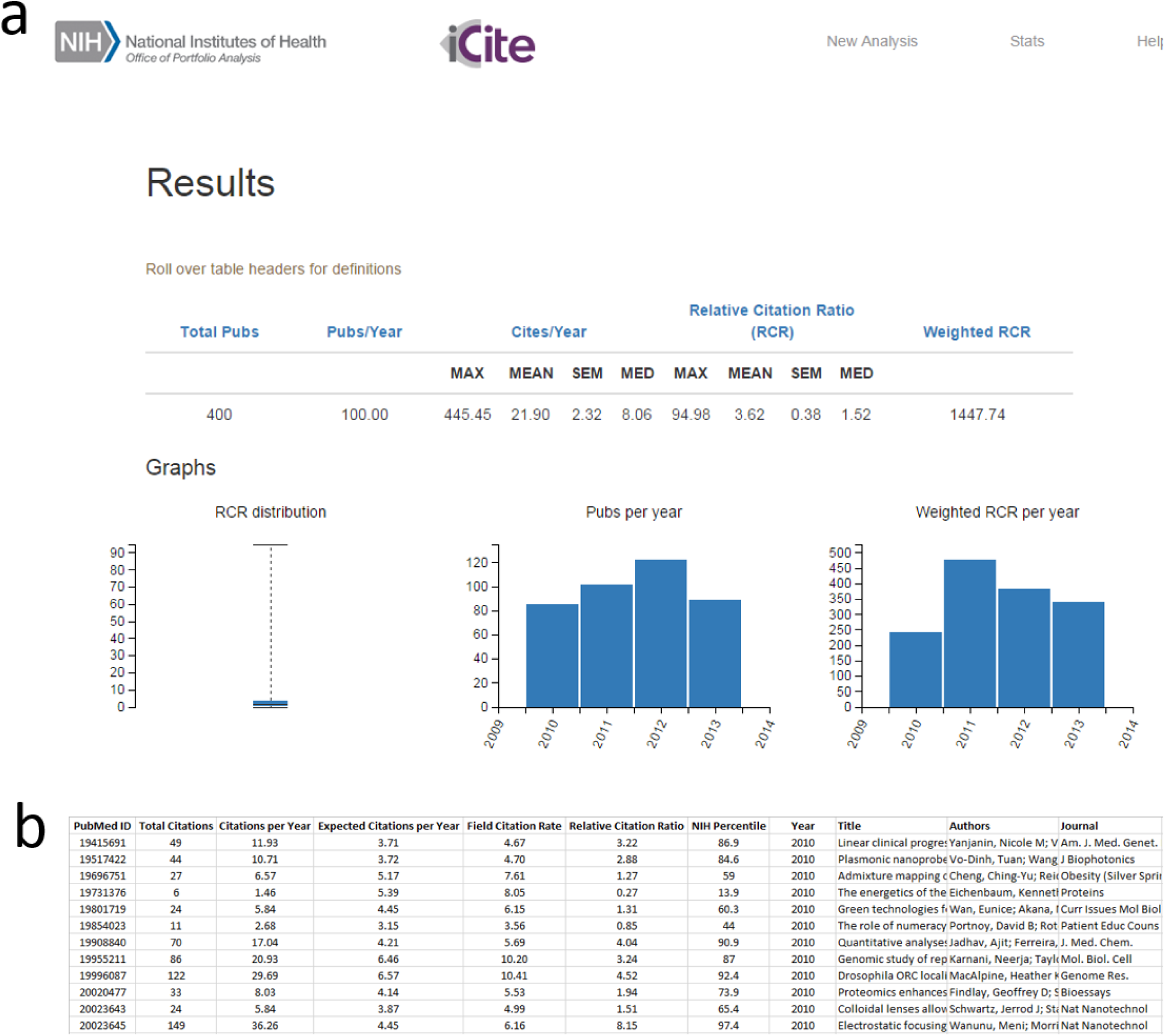
*iCite*, a publicly available tool for calculating RCR and accessing related citation information. **(a)** Screen shot of a sample *iCite* result. 400 sample PMIDs from papers published over a four year window were entered into the *iCite* tool, which returned the maximum, mean */- standard error of the mean (SEM), and median values for both citations per year and RCR; weighted RCR is equal to the sum of the RCRs for this group. Box and whisker shoes the distribution of article RCRs; bar graphs show the number of publications per year and weighted RCR per year, respectively. **(b)** Sample data download for an *iCite* result. *iCite* returns the total number of citations, number of citations per year, expected citations per year based on an NIH R01 benchmark, field citation rate, relative citation rate, and percentile ranking in a downloadable Excel format for each PMID entered, as well as the corresponding title, author information, and year/journal of publication.

### RCR-based evaluation of two NIH-funded research programs

One of the unique strengths of RCR is the way in which a paper’s co-citation network dynamically defines its field. Each new citation an article receives, then, can be thought of as originating either from within or from outside its existing network. As a work gains relevance to additional disciplines, it seems intuitively possible that a new out-of-network citation might lead to a disproportionate increase in Field Citation Rate and thus to a drop in RCR. Such an occurrence might be thought undesirable (40); alternatively, it might be considered an accurate reflection of the reduced relative influence the work in question has on a new and larger group of scholars, many of whom previously may not have had a reason to encounter it. Regardless, we felt it was important to determine how frequently such a hypothetical scenario (40) occurs. Among the more than 200,000 articles published between 2003 and 2011 for which we calculated Field Citation Rates, less than 2% experienced any sort of drop in RCR between 2012 and 2014; only 0.2% experienced a drop in RCR of 0.1 or more. This low incidence is consistent with the stability we observe in Field Citation Rates (**Figure 3c**) and with the theoretical properties of citation networks, which are known to be scale-free and thus resistant to perturbation (48).

While 0.2% is a very small number, we wondered whether interdisciplinary science might be overrepresented among the articles that did experience a drop in RCR. An impediment to testing this hypothesis, though, is the lack of a precisely circumscribed consensus definition of interdisciplinary research; one reason it is difficult to arrive at such a definition is that disciplines are themselves dynamic and undergo continuous evolution. For example, biochemistry may have been considered highly interdisciplinary in 1905, the year that term first appears in the PubMed indexed literature (49), but most biomedical researchers today would consider it

### RCR-based evaluation of two NIH-funded research programs

One of the unique strengths of RCR is the way in which a paper’s co-citation network dynamically defines its field. Each new citation an article receives, then, can be thought of as originating either from within or from outside its existing network. As a work gains relevance to additional disciplines, it seems intuitively possible that a new out-of-network citation might lead to a disproportionate increase in Field Citation Rate and thus to a drop in RCR. Such an occurrence might be thought undesirable (40); alternatively, it might be considered an accurate reflection of the reduced relative influence the work in question has on a new and larger group of scholars, many of whom previously may not have had a reason to encounter it. Regardless, we felt it was important to determine how frequently such a hypothetical scenario (40) occurs. Among the more than 200,000 articles published between 2003 and 2011 for which we calculated Field Citation Rates, less than 2% experienced any sort of drop in RCR between 2012 and 2014; only 0.2% experienced a drop in RCR of 0.1 or more. This low incidence is consistent with the stability we observe in Field Citation Rates (**Figure 3c**) and with the theoretical properties of citation networks, which are known to be scale-free and thus resistant to perturbation (48).

While 0.2% is a very small number, we wondered whether interdisciplinary science might be overrepresented among the articles that did experience a drop in RCR. An impediment to testing this hypothesis, though, is the lack of a precisely circumscribed consensus definition of interdisciplinary research; one reason it is difficult to arrive at such a definition is that disciplines are themselves dynamic and undergo continuous evolution. For example, biochemistry may have been considered highly interdisciplinary in 1905, the year that term first appears in the PubMed indexed literature (49), but most biomedical researchers today would consider it a well-established discipline in its own right. Others might still view it as an interdisciplinary field in the strictest sense, as it occupies a space between the broader fields of biology and chemistry. To some extent, then, interdisciplinarity is in the eye of the beholder, and this presents another challenge. The question only becomes more vexed when considering more recent mergers such as computational biology, neurophysiology, or developmental genetics; are these established fields, interdisciplinary fields, or sub-fields? As a first approximation, then, we chose to ask whether articles produced by the NIH Interdisciplinary Research Common Fund program, which funded work that conforms to the definition of interdisciplinarity adopted by the National Academy of Sciences (50,51), were more or less likely to experience a drop in RCR than other NIH funded articles. Interestingly, these interdisciplinary papers were actually two-fold less likely to experience a drop in RCR than papers funded either by other Common Fund programs or by standard R01 support (**Figure 6a**). We currently lack an explanation as to why this might be so; given how infrequent these small drops are, we cannot yet rule out the possibility that statistical noise is responsible.

**Figure 6.**
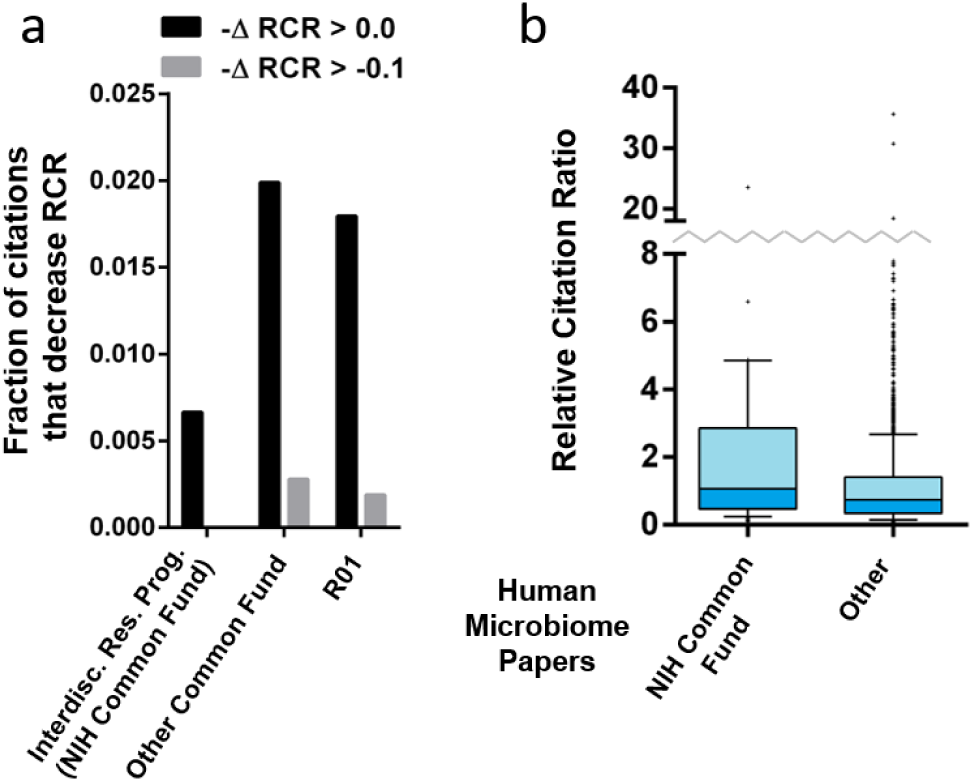
RCR-based evaluation of two NIH-funded research programs. **(a)** Bar graph showing the percentage of papers that experience a drop in RCR from 2012 to 2014. Black bars, any decrease in RCR; gray bars, decrease in RCR of 0.1 or more. **(b)** Box-and-whisker plots showing the distribution of RCR values for articles describing the human microbiome, published with support from the Human Microbiome Project of the NIH Common Fund (HMP) or another source (other). Boxes show the 25th-75th percentiles with a line at the median; whiskers extend to the 10th and 90th percentiles.

We also analyzed publications funded by NIH’s Human Microbiome Project (HMP), which was established in 2007 to generate research resources to facilitate the characterization and analysis of human microbiota in health and disease. From 2009-2011, scientists funded by the HMP published 87 articles for which citation information is available. As a comparison group, we identified 2267 articles on the human microbiome that were published during the same time period, but were not funded by the HMP. Articles from the Human Microbiome Project outperformed the comparison group (**Figure 6b**; HMP mean RCR 2.30, median RCR 1.06; comparison RCR mean 1.23, median 0.74; p < 0.001, Mann-Whitney U test), demonstrating that sorting by funding mechanism has the potential to identify works of differential influence.

### Quantifying how past influence predicts future performance

We next undertook a large case study of all 88,835 articles published by NIH investigators who maintained continuous R01 funding from fiscal year (FY) 2003 through FY2010 to ask how the RCR of publications from individual investigators changed over this eight-year interval. Each of these investigators had succeeded at least once in renewing one or more of their projects through the NIH competitive peer review process. In aggregate, the RCR values for these articles are well-matched to a log-normal distribution; in contrast, as noted previously by others, the distribution of impact factors of the journals in which they were published is non-normal (52,53) (**Figure 7a**, **b**). Sorting into quintiles based on JIF demonstrates that, though journals with the highest impact factors have the highest median RCR, influential publications can be found in virtually all journals (**Figure 7c**, **d**). Focusing on a dozen representative journals with a wide range of JIFs further substantiates the finding that influential science appears in many venues, and reveals noteworthy departures from the correlation between JIF and median RCR (see **Supporting Information**). For example, NIH-funded articles in both *Organic Letters* (JIF = 4.7) and the *Journal of the Acoustical Society of America* (JIF = 1.6) have a higher median RCR than those in *Nucleic Acids Research* (JIF = 7.1; **Figure 7e**).

**Figure 7.**
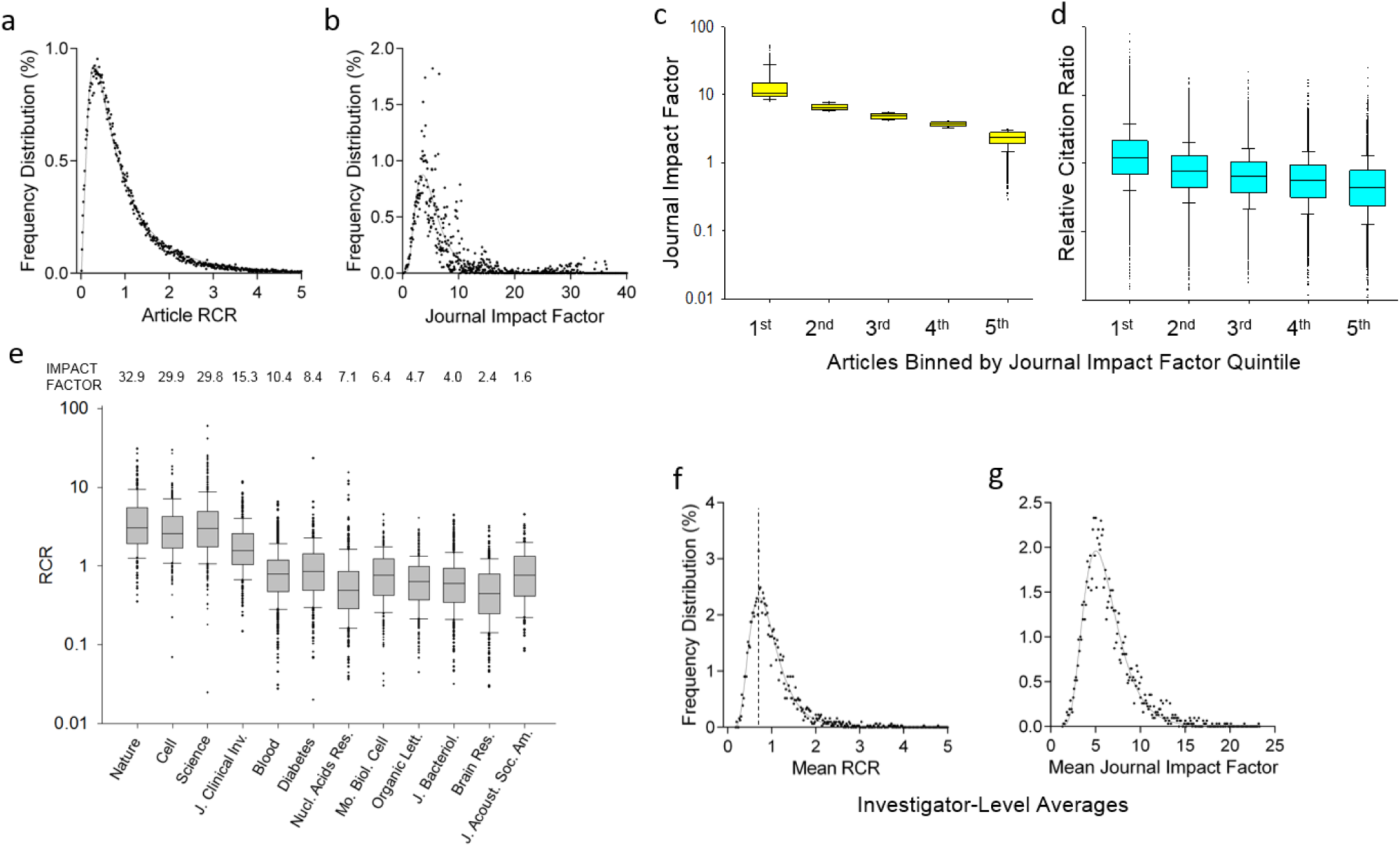
Properties of Relative Citation Ratios at the article and investigator level. (**a, b**) Frequency distribution of article-level RCRs (**a**) and Journal Impact Factors (**b**), from 88,835 papers (authored by 3089 R01-funded PIs) for which co-citation networks were generated. Article RCRs are well-fit by a log-normal distribution (R^2^ = 0.99), and Journal Impact Factors less so (R^2^ = 0.79). (**c**) Box-and-whisker plots summarizing Journal Impact Factors for the same papers, binned by Impact Factor quintile (line, median; box, 25th–75th percentiles; whiskers, 10th to 90th percentiles). (**d**) RCR for the same papers using the same bins by Journal Impact Factor quintile (same scale as c). Although the median RCR for each bin generally corresponds to the Impact Factor quintile, there is a wide range of article RCRs in each category. (**e**) Box-and-whisker plots summarizing RCRs of these same papers published in selected journals. In each journal, there are papers with article RCRs surpassing the median RCR of the highest Impact Factor journals (left three). The Impact Factor of each journal is shown above. (**f, g**) Frequency distribution of investigator-level RCRs (**f**) and Journal Impact Factors (**g**), representing the mean values for papers authored by each of 3089 R01-funded PIs. Dashed line in (**f**), mode of RCR for PIs.

As part of this case study we also calculated the average RCR and average JIF for papers published by each of the 3089 NIH R01 principal investigators (PIs) represented in the dataset of 88,835 articles. In aggregate, the average RCR and JIF values for NIH R01 PIs exhibited log-normal distributions (**Figure 7f**, **g**) with substantially different hierarchical ordering (**Supporting Figure S8**). This raised a further question concerning PIs with RCR values near the mode of the log-normal distribution (dashed line in **Figure 7f**): as measured by the ability to publish work that influences their respective fields, to what extent does their performance fluctuate? We addressed this question by dividing the eight year window (FY2003 through FY2010) in half. Average RCRs in the first time period (FY2003 through FY2006) were sorted into quintiles, and the percentage of PIs in the second time period (FY2007 through FY2010) that remained in the same quintile, or moved to a higher or lower quintile, was calculated. The position of PIs in these quintiles appeared to be relatively immobile; 53% of PIs in the top quintile remained at the top, and 53% of those in the bottom quintile remained at the bottom (**Figure 8a**). For each PI we also calculated a weighted RCR (the number of articles multiplied by their average RCR); comparing on this basis yielded almost identical results (**Figure 8b**). It is worth noting that average Field Citation Rates for investigators were extremely stable from one 4-year period to the next (Pearson ***r*** = 0.92, **Table 2**), Since Field Citation Rates are the quantitative representation of cocitation networks, this further suggests that each cocitation network is successfully capturing the corresponding investigator’s field of research.

**Figure 8.**
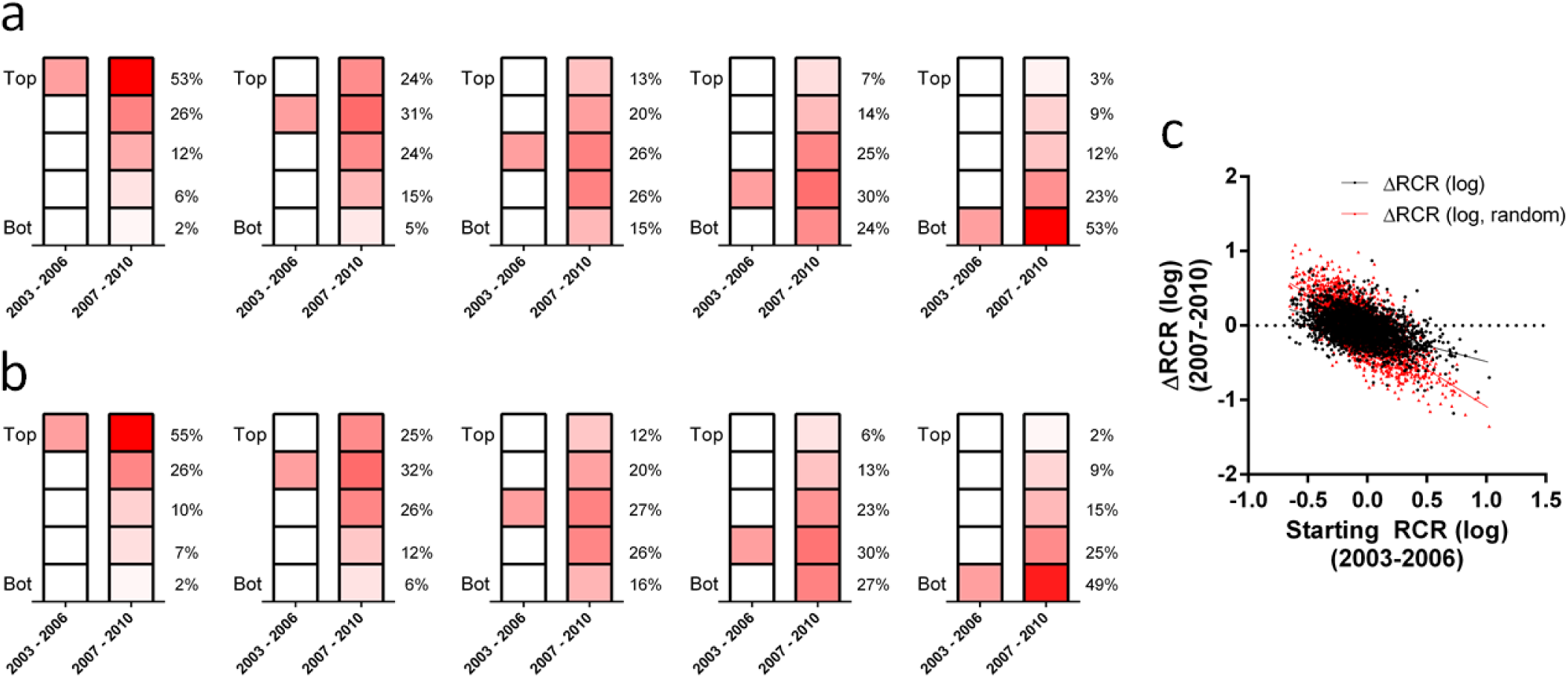
Scientific mobility of investigators’ influence relative to their field. Color intensity is proportional to the percentage of PIs in each quintile. (**a**) 3089 investigators who were continuously funded by at least one R01 were ranked by their articles’ average RCR in each time window, and split into quintiles. From left to right, investigators starting in different quintiles were tracked to see their rank in the next 4-year period. (**b**) The same analysis, but the number of published articles was multiplied by their average RCR to calculate an influence-weighted article count. PIs were ranked by this aggregate score and split into quintiles. (**c**) Scatter plot illustrating the relationship between PI RCR at earlier and later time frames. Black points, actual RCR values; black line, linear regression of actual RCR values. Red points, random assignment model (PI RCRs for the second four year period are re-shuffled and randomly assigned); red line, linear regression of modeled data.

**Table 2.** Summary of investigator-level bibliometric measures and their stability from one 4-year period to the next (PIs with more than 5 articles in each period, except for article count).

| Measure | Mean | Median | Pearson *r*<br>(of log-values,<br>'03-'06 vs '07-'10) |
| --- | --- | --- | --- |
| Articles per interval<br>(PIs with >0 articles in<br>both 4-yr. intervals) | 9.8 | 8.0 | 0.56 |
| Field citation rate | 7.8 | 7.7 | 0.92 |
| Journal citation rate | 6.3 | 5.7 | 0.76 |
| Article citation rate | 6.4 | 5.3 | 0.67 |
| RCR | 1.0 | 0.85 | 0.61 |

Another possible interpretation of the above data is that PI RCRs perform an unbiased random walk from their initial state with a large diffusion rate. Considered from this frame, it could be said that 47% of PIs who started in the top quintile moved out of it during the second 4-year period we analyzed. To test this hypothesis directly, we performed a mean reversion test, which determines whether or not the set of values under consideration will return to an average, or mean, value over time. If drift in PI RCR were simply a random walk, then the change in RCR should by definition be independent of starting RCR, and plotting these two values against each other should result in a straight line with a slope of zero. However, the results show that change in RCR is dependent on starting RCR value (p < 0.001, linear regression analysis, n = 3089, **Figure 8c**). Furthermore, randomly shuffling PI RCRs from the second four year period gives a slope that is significantly different than that observed for the real data (p < 0.001, Extra sum-of-squares F-test, n = 3089, **Figure 8c**), ruling out the possibility that these values are randomly sampled from the same distribution in each time interval.

## Discussion

The relationship between scientists and JIFs has been likened to the prisoner’s dilemma from game theory: because grant reviewers use JIFs in their evaluations, investigators must continue to weigh this in their decision-making or risk being out-competed by their peers on this basis (54,55). A groundswell of support for the San Francisco Declaration on Research Assessment (http://www.ascb.org/dora) has not yet been sufficient to break this cycle (54–59). Continued use of the Journal Impact Factor as an evaluation metric will fail to credit researchers for publishing highly influential work. Articles in high-profile journals have average RCRs of approximately 3. However, high-Impact-Factor journals (JIF ≥ 28) only account for 11% of papers that have an RCR of 3 or above. Using Impact Factors to credit influential work therefore means overlooking 89% of similarly influential papers published in less prestigious venues.

Bibliometrics like JIF and H-index are attractive because citations are affirmations of the spread of knowledge amongst publishing scientists, and are important indicators of the influence of a particular set of ideas. Though tracking the productivity of individual scientists with bibliometrics has been controversial, it is difficult to contradict the assertion that uncited articles (RCR = 0) have little if any influence on their respective fields, or that the best-cited articles (RCR > 20) are impressively influential. We have not determined whether smaller differences, for example those with average or slightly above-average RCRs (e.g. 1.0 versus 1.2), reliably reflect differential levels of influence. Further, citation-based metrics can never fully capture all of the relevant information about an article, such as the underlying value of a study or the importance of making progress in solving the problem being addressed. The RCR metric is also not designed to be an indicator of longterm impact, and citation metrics are not appropriate for applied research, e.g. work that is intended to target a narrow audience of non-academic engineers or clinicians.

It is also very important to note that like all other citation-based metrics, an RCR value cannot be calculated immediately after an article is published. Instead, enough time must pass for a meaningful number of citations to accrue, and the work we describe here provides some rough guidance as to what that meaningful number might be. Specifically, we have found that 93% of co-citation network-based field citation rates stabilize after a work has been cited five times (**Figure 3c**); also in agreement with previously published work, citation rates for approximately the same percentage of articles peak within two to three years after publication (**Supporting Figure S2**). Before one or both of those benchmarks have been reached, RCR values might be viewed as provisional; even after that point, neither RCR nor any other citation-based metric should be taken as a substitute for the actual reading of a paper in determining its quality. However, as citation rates mark the breadth and speed of the diffusion of knowledge among publishing scholars, we maintain that quantitative metrics based on citations can effectively supplement subject matter expertise in the evaluation of research groups seeking to make new discoveries and widely disseminate their findings.

We believe RCR offers some significant advantages over existing citation-based metrics, both technically and in terms of usability. Technically, prior attempts to describe a normalized citation metric have resulted in imperfect systems for the comparison of diverse scholarly works (**Supporting Information** and **Figure 4e**), either because they measure only the average performance of a group of papers (60), or because the article of interest is measured against a control group that includes widely varying areas of science (17,31,47). An example of the latter is citation percentiling, which the Leiden manifesto (42) recently recommended as best practice in bibliometrics. Theoretically, the RCR method is an improvement over the use of citation percentiling alone, since masking the skewed distribution of citations and article influence, while statistically convenient, can disadvantage portfolios of high-risk, high-reward research that would be expected to have a small proportion of highly influential articles (61). Furthermore, we have shown that co-citation networks better define an article’s field than journal of publication (**Figure 2**), so RCR is a more precise measure of influence than journal-based metrics, a category that includes both citation percentiles and MNCS methods such as the TR ratio. RCR is also no more likely to unfairly advantage publications in fields with a low citation rate than the TR ratio (**Figure 4e**). Finally, by incorporating a way to benchmark to a meaningful comparison group, RCR makes it easy for users to know whether a set of articles is above or below expectations for their region or agency; the need for such a feature has already been prominently discussed (30).

In terms of usability, both RCR values and their component variables, including Field Citation Rates, citations per year, and total citations are freely available to the public through our *iCite* tool. As much of the source citation data is proprietary, we are prevented from identify all of the citing papers; presently, all bibliometrics face this challenge, as limited open source citation data is available. We feel that RCR and *iCite* represent a large improvement in transparency relative to citation percentiles and TR ratios, which are not cost-free, and are furthermore dependent on the proprietary classification of journals into one or another area of science. Our method and tool are also far more transparent than impact factor, the calculation of which has recently come under scrutiny after allegations of manipulation (56,62,63).

Any metric can be gamed, and we have thought carefully about how a single author might try to game RCR. Article citation rates could be inflated through a combination of self-citation and frequent publication; this strategy has its limits, though, as the top 10% of RCR values for NIH-funded publications on average receive more than 25 citations per year, and it is rare for a biomedical scientist to publish more than four or five times over that period. A more promising strategy might be to strive for the lowest possible Field Citation Rate. An author taking this approach would need to stack the reference section of his or her work not just with poorly cited articles, or with articles in poorly cited fields, but with articles that are co-cited with articles in poorly cited fields. Since citing behavior is also constrained by content, this might be difficult to accomplish; at the very least, it seems likely that reviewers and editors would be able to identify the resulting reference list as unusual. Of course, if enough authors start to reference works in poorly cited areas, that field’s citation rate will go up, and the RCR of the papers in it may go down; in that respect, efforts to game RCR might ultimately prove to be self-defeating.

An important point to keep in mind when interpreting RCR values, though, is that citations follow a power law or log-normal distribution (9), wherein one researcher’s choice of a particular reference article is at least partly informed by the choices that other researchers have previously made. There is a certain amount of noise inherent in that selection process (26), especially in the early days of a new discovery when a field is actively working towards consensus. The results of a landmark study on the relationship between quality and success in a competitive market suggests that the ultimate winners in such contests are determined not only by the intrinsic value of the work, but also by more intangible social variables (64). Consistent with this conclusion, a different group of authors have shown that including a “reputation” variable enables an algorithm to better predict which papers in the interdisciplinary field of econophysics will be the most highly cited (65). Although there is on average a positive relationship between quality and success (51), it is for this reason we suggest that RCR should primarily be considered as a measure of influence, rather than impact or intellectual rigor.

Within these bounds, bibliometric methods such as RCR have the potential to track patterns of scientific productivity over time, which may help answer important questions about how science progresses. In particular, co-citation networks can be used to characterize the relationship between scientific topics (including interdisciplinarity), emerging areas, and social interactions. For example, is the membership of an influential group of investigators in a given field or group of fields stable over time, or is it dynamic, and why? Our data demonstrate the existence of an established hierarchy of influence within the exclusive cohort of NIH R01 recipients who remained continuously funded over an eight-year time frame. This may mean that investigators tend to ask and answer questions of similar interest to their fields. Additionally or alternatively, stable differences in investigators’ status, such as scientific pedigree, institutional resources, and/or peer networks, may be significant drivers of persistently higher or lower RCR values. Future statistical analyses may therefore reveal parameters that contribute to scholarly influence. To the extent that scientific (im)mobility is a product of uneven opportunities afforded to investigators, there may be practical ways in which funding agencies can make policy changes that increase mobility and seed breakthroughs more widely.

There is increasing interest from the public in the outcomes of research. It is therefore becoming necessary to demonstrate outcomes at all levels of funding entities’ research portfolios, beyond the reporting of success stories that can be quickly and succinctly communicated. For this reason, quantitative metrics are likely to become more prominent in research evaluation, especially in large-scale program and policy evaluations. Questions about how to advance science most effectively within the constraints of limited funding require that we apply scientific approaches to determine how science is funded (66–69). Since quantitative analysis will likely play an increasingly prominent role going forward, it is critical that the scientific community accept only approaches and metrics that are demonstrably valid, vetted, and transparent, and insist on their use only in a broader context that includes interpretation by subject matter experts.

Recent work has improved our theoretical understanding of citation dynamics (26–28). However, citation counts are not the primary interest of funding agencies, but rather progress in solving scientific challenges. The NIH particularly values work that ultimately culminates in advances to human health, a process that has historically taken decades (70). Here too, metrics have facilitated quantitation of the diffusion of knowledge from basic research toward human health studies, by examining the *type* rather than the *count* of citing articles (71). Insights into how to accelerate this process will probably come from quantitative analysis. To credit the impact of research that may currently be underappreciated, comprehensive evaluation of funding outputs will need to incorporate metrics that can capture many other types of outputs, outcomes, and impact, such as the value of innovation, clinical outcomes, new software, patents, and economic activity. As such, the metric described here should not be viewed as a tool to be used as a primary criterion in funding decisions, but as one of several metrics that can provide assistance to decision-makers at funding agencies or in other situations in which quantitation can be used judiciously to supplement, not substitute for, expert opinion.

## Materials and Methods

### Citation data

The Thomson Reuters Web of Science citation dataset from 2002-2012 was used for citation analyses. For FCR stability analysis this dataset was extended to include 2014 data. Because of our primary interest in biomedical research, we limited our analysis to those journals in which NIH R01-funded researchers published during this time. For assigning a journal citation rate to a published article, we used the 2-year synchronous journal citation rate (38,72) for its journal in the year of its publication. Publications from the final year of our dataset (2012) were not included in analyses because they did not have time to accrue enough citations from which to draw meaningful conclusions, but references from these papers to earlier ones were included in citation counts.

### Grant and Principal Investigator data

Grant data was downloaded from the NIH RePORTER database (https://proiectreporter.nih.gov/). Grant-to-publication linkages were first derived from the NIH SPIRES database, and the data were cleaned to address false-positives and -negatives. Grant and publication linkages to Principal Investigators were established using Person Profile IDs from the NIH IMPAC-II database. To generate a list of continuously funded investigators, only those Person Profile IDs with active R01 support in each of Fiscal Years 2003-2010 were included.

### Calculations and data visualization

Co-citation networks were generated in Python (Python Software Foundation, Beaverton, OR). This was accomplished on a paper-by-paper basis by assembling the list of articles citing the article of interest, and then assembling a list of each paper that those cited. This list of co-cited papers was de-duplicated at this point. Example code for generating co-citation networks and calculating Field Citation Rates is available on GitHub (http://github.com/NIHOPA). Data that was used for analysis can be found as csv files in the same repository. Further calculations were handled in R (R Foundation for Statistical Computing, Vienna, Austria). Visualizations were generated in Prism 6 (GraphPad, La Jolla, CA), SigmaPlot (Systat Software, San Jose, CA), or Excel 2010 (Microsoft, Redmond, WA). Code used to generate the database used in the *iCite* web application (https://icite.od.nih.gov) can be found in the GitHub repository. A preprint version of this manuscript can also be found on bioRxiv (73). For box-and-whisker plots, boxes represent the interquartile range with a line in between at the median, and whiskers extend to the 10^th^ and 90^th^ percentiles.

When comparing citations rates to other metrics (e.g. post-publication review scores), citation rates were log-transformed due to their highly skewed distribution, unless these other scores were similarly skewed (i.e. Faculty of 1000 review scores). For this process, article RCRs of zero were converted to the first power of 10 lower than the lowest positive number in the dataset (generally 10^−2^). In the analysis of Principal Investigator RCRs, no investigators had an average RCR of zero.

### Content analysis

The commercially available text mining program IN-SPIRE (Pacific Northwest National Laboratories, Richland, WA; [74]) was used for content-based clustering of papers in a co-citation network (**Figure 1c**). For comparison of journal impact factor citations per year and RCR (**Figure 4d**), papers in the fields of cell biology and neurological function were those supported by grants assigned to the corresponding peer review units within the NIH Center for Scientific Review.

For the data in **Figure 2**, articles were selected from six journals; three of these were disciplinary (Journal of Neuroscience, Blood, and Genetics) and the other three were multidisciplinary (Nature, Science and PNAS). Articles with exactly five citations were chosen to limit the number of pairwise comparisons in the co-citation network. Abstracts (from PubMed) were concatenated with titles to comprise the information for each document. Words were converted to lower case and stemmed. Any numbers, as well as words consisting of one or two letters, were removed from the corpus along with words appearing less than ten times. Term-document matrices were weighted for term frequency-inverse document frequency (TF-IDF) (75). For one analysis the term-document matrix was trimmed to the top 1000 TF-IDF-weighted terms, and in the other analysis, no additional term trimming was performed. Cosine similarity scores (36) were calculated for 1397 reference articles against each article in their co-citation network, and separately against each article appearing in the same journal. This resulted in 249,981 pairwise comparisons with articles in the co-citation networks, and 28,516,576 pairwise comparisons with articles from the same journals. Frequency distributions for each journal and the co-citation network comparison are shown in **Figure 2**.

## Acknowledgements

We thank Francis Collins, Larry Tabak, Kristine Willis, Mike Lauer, Jon Lorsch, Stefano Bertuzzi, Steve Leicht, Stefan Maas, Riq Parra, Dashun Wang, Patricia Forcinito, Carole Christian, Adam Apostoli, Aviva Litovitz and Paula Fearon for their thoughtful comments on the manuscript, Michael Gottesman for help organizing postpublication peer review, and Jason Palmer, Fai Chan, Rob Harriman, Kirk Baker and Kevin Small for help with data processing and software development for *iCite*.

## Supporting Information

### Characterization of the co-citation networks of single articles

Field normalization of citations is critical for cross-field comparisons, because of intrinsic differences in citation rates across disciplines. Our approach draws upon the idea of comparing the Article Citation Rate of an article (ACR) with an Expected Citation Rate (ECR), calculated based on peer performance in an article’s area of research (1,2). Calculating a robust ECR is challenging; other methods frequently employ journals or journal categories as a proxy for a scientific field (3–9). Unfortunately, these methods do not have sufficient precision to work well at the article level (10). Because modern fields of biomedical research exist as a spectrum rather than discrete and separate fields (11), we decided to take a more nuanced approach to defining an article’s field. We constructed each article’s co-citation network (12) and used that as a representative sample of its area of research. Simply put, when an article is first cited, the other papers appearing in the reference list along with the article comprise its co-citation network (**Figure 1**). As the article continues to be cited, the papers appearing in the new reference lists alongside it are added to its co-citation network. This network provides a dynamic view of the article’s field of research, taking advantage of information provided by the experts who have found the study useful enough to cite.

### Algorithm and calculations for Relative Citation Ratios (RCRs)

Our algorithm uses the following steps to calculate RCR values, giving a ratio of ACR to ECR that is benchmarked to papers funded through NIH R01s:

1. Convert the RA citation counts to citations per year (**Figure 3** and **Supporting Equation S1**).
2. Generate the RA’s co-citation network. To do this, we assemble all articles citing the RA; the complete set of papers cited in the reference lists of these citing articles comprises the co-citation network (**Figure 1**).
3. Estimate the FCR of the RA by averaging the journal citation rates of the papers in the co-citation network (**Supporting Equation S3**).
4. Generate an ECR from the benchmark set of papers. Using R01-funded papers published in a given year, a linear regression of the ACRs vs. FCRs is performed (**Figure 2** and **Supporting Equations S3–4**). Regressions are calculated for each publication year.

a. The linear equation coefficients corresponding to the RA’s publication year rescale its FCR into a denominator (ECR) that is benchmarked to the performance of R01-funded articles (**Supporting Equation S5**).
5. The Relative Citation Ratio is the ratio of the ACR : ECR.

When converting raw citation counts to Article Citation Rate (*Acr*), the year in which the RA was published was excluded from the denominator.

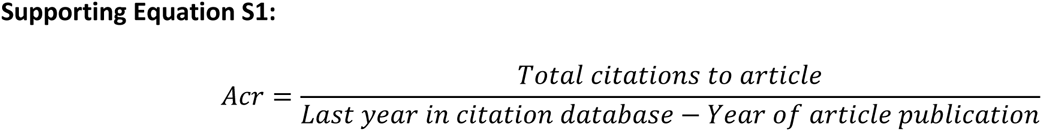

We made this design decision because the publication year is nearly always partial, and because articles receive a low number of citations in the calendar year of their publication compared to subsequent years (**Supporting Figure S2**). In practice, the sum of the citations in years 0 and 1 (the year of publication and the following year) is close to the mean number of citations per year in the following 8 years (**Supporting Figure S2**).

ACRs and journal citation rates vary widely from field to field. To compare the Relative Citation Ratios of small groups (like individual investigators), special care must be taken to adjust only the fraction of the citation rate that is due to between-field differences. We tested three methods for adjusting expected citation rates to a field. These approaches each use information from an article’s citation network (see schematics in **Figure 1a**). The first method selects the reference article plus those cited in in its reference list (**Figure 1a**, bottom). The second approach instead selects subsequent papers citing the article (**Figure 1a**, top). Finally, the third uses the set of articles that are co-cited with the article of interest by subsequent papers (**Figure 1a**, middle). The Reference Article was always included in the set of papers selected to estimate the FCR, since an article is de facto part of its field. In all three cases, the average of the journal citation rates for the papers in the selected level of the citation network is used as the Field Citation Rate (*Fcr*).

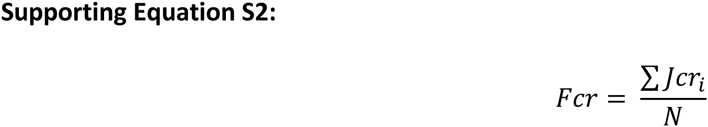

*N* is the number of papers in the selected level of the co-citation network (**Figure 1a**) and *Jcr_i_* is the journal citation rate of each paper at the specified level of the citation network.

To generate an expectation of citation performance using this cohort of papers, we performed a linear regression of Article Citation Rate (*Acr*) in a baseline population against their Field Citation Rates (*Fcr*) from the same year (**Figure 3d**). R01-funded articles were used as a benchmark population. This process was repeated for each year being analyzed to give regression coefficients (slope, *B̂* and intercept, *â*) for benchmarking articles.

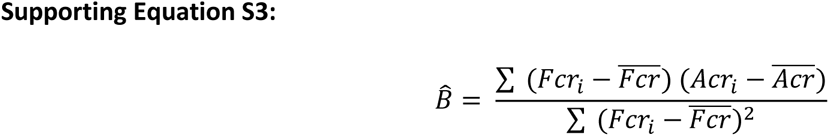

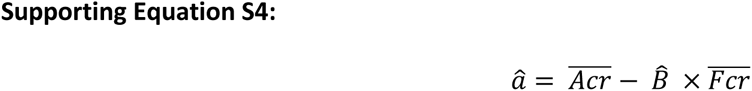

Alternatively, if benchmarking to the median (rather than mean) field-normalized performance for the benchmark group is desired, quantile regression can be used in lieu of simple linear regression (13). In either case, the resulting regression line transforms a FCR into an Expected Citation Rate (*Ecr^Year^*), corresponding to the ACR that R01-funded papers with the same FCRs and published in the same year were able to achieve, and this can be used outside of the baseline population as a benchmark for articles published in that year:

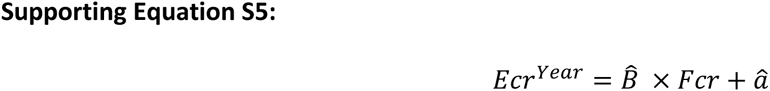

For the years analyzed here (2002–2011), the resulting regression coefficients are given in Supporting Table S2. RCR for each article is the ratio of that article’s ACR divided by its ECR. Calculating the arithmetic mean is the preferred way of determining the RCR of an entire portfolio (6–8):

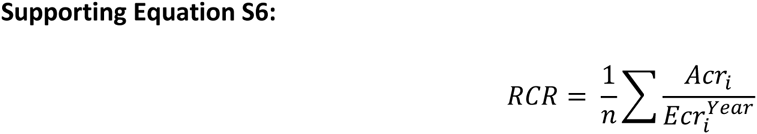

Where *n* is the number of papers being evaluated, *Acr_i_* is the article citation rate and 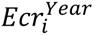 the expected citation rate of each article found by transforming its *Fcr_i_* with the regression coefficients for its publication year (**Supporting Equation S5**).

### Evaluation of different levels of the citation network for field normalization

The aim of this adjustment is to accurately normalize an individual article’s citation rate to its field’s average citation rate, while preserving within-field differences between papers. Using a set of papers from 2009 matched to R01 grants active in the same year, we compared these three approaches. Calculated RCRs were very similar for all three groups at the article level (**Supporting Table S3**). This is not surprising, since at this granular scale the article’s numerator (ACR) accounts for more of the variance than the denominator. However, accurate field-adjustment is especially important for large-scale analyses, where field differences in citation rates can dominate measurements, as article-level differences in ACR average out. A more accurate estimate of the field citation rate would be predicted to show a smaller correlation between article citation rate and field citation rate, as within-field differences are more effectively excluded. To measure the effectiveness of each level of the citation network, we calculated the correlations between ACRs and ECRs for each approach. Of the three, the “Co-cited” method shows the least correlation between article citations and expected citations (**Supporting Table S4**). In addition, the variance in expected citation rates should be lower in approaches that more successfully isolate the between-field differences in citation rate from the within-field differences. Again, the co-citation level of the citation network performed the best here (**Table 1**).

### Validation of RCR with post-publication peer review

We extensively validated article-level RCRs against expert reviewer scores of the impact or value of papers, using post-publication peer review. Three independently collected sets of post-publication peer reviews were used for this analysis: Faculty of 1000 (F1000) (5,14), a previous survey conducted by the Institute for Defense Analyses Science and Technology Policy Institute (STPI) (15), and post-publication peer review conducted by NIH Intramural Research Program (IRP) Principal Investigators.

The first set of expert review scores was compiled from F1000, in which faculty review articles in their fields of expertise, and rate the articles on a scale of 1 to 3 (“Good”, “Very Good”, and “Exceptional”). Because the decision by the faculty members to review the article is itself a mark of merit, these scores are summed into a composite F1000 score (**Supporting Figure S3**). We downloaded scores in June 2014 for 2193 R01-funded articles published in 2009 and compared them to their RCRs. This yielded an article-level correlation coefficient ***r*** of 0.44 between RCR and F1000 scores (**Figure 4a**).

For a second set of expert review scores, we took advantage of a previous survey conducted by STPI, of papers funded through the Howard Hughes Medical Institute and NIH. In this survey, experts rated the impact of articles (shown here on a scale of 0 to 4, n = 430 papers from 2005-2011, **Supporting Figure S4**). Since citation data (including RCR) is highly skewed while survey ratings are not, RCR was log-transformed to bring these ranges into better alignment. The article-level correspondence of RCR with these review scores was similar to that observed with the F1000 scores (***r*** = 0.47, **Figure 4b**).

Finally, we recruited investigators from the NIH Intramural Research Program (IRP) to perform postpublication peer review of NIH-funded articles published in 2009 (**Supporting Figures S5–S7**). A total of 290 articles were independently reviewed by multiple investigators, yielding an article-level correlation of 0.56 (**Figure 4c**). The distribution of these impact scores is shown in Figure 2 Supplement 8. Finally, we asked reviewers in the NIH Intramural Research Program to conduct post-publication peer review of R01-funded articles published in 2009 (reviews conducted by The Scientific Consulting Group, Gaithersburg, MD). Reviewers were asked to give scores on a scale of 1-5 for the following questions:

- Rate whether the question being addressed is important to answer. (1 = Not Important, 2 = Slightly Important, 3 = Important, 4 = Highly Important, 5 = Extremely Important)
- Rate whether you agree that the methods are appropriate and the scope of the experiments adequate. (1 = Strongly Disagree, 2 = Disagree, 3 = Neutral, 4 = Agree, 5 = Strongly Agree)
- Rate how robust the study is based on the strength of the evidence presented. (1 = Not Robust, 2 = Slightly Robust, 3 = Moderately Robust, 4 = Highly Robust, 5 = Extremely Robust)
- Rate the likelihood that the results could ultimately have a substantial positive impact on human health outcomes. (1 = Very unlikely, 2 = Unlikely, 3 = Foreseeable but uncertain, 4 = Probable, 5 = Almost Certainly)
- Rate the impact that the research is likely to have or has already had. (1 = Minimal Impact, 2 = Some Impact, 3 = Moderate Impact, 4 = High Impact, 5 = Extremely High Impact)
- Provide your overall evaluation of the value and impact of this publication. (1 = minimal or no value, 2 = Moderate value, 3 = Average value, 4 = High value, 5= Extremely high value)

The distribution of responses for each of these questions is shown in **Supporting Figure S5**. Multiple experts were asked to review each paper, and each set of papers were matched to the fields of expertise of the reviewers examining them. For correlating RCR to review scores, we used the average of the score for the final question (overall evaluation of the paper’s value) for articles that were reviewed by at least two experts.

To determine post-hoc which of the first five criteria (importance of scientific question, appropriate methods, robustness of study, likelihood of health outcomes and likely impact) were associated with reviewers’ ratings of overall value, we first performed a Random Forest analysis using all 5 criteria. Perhaps unsurprisingly, this analysis showed that “likely impact” was most closely associated with assessment of overall value (**Supporting Figure S6a**). We subsequently removed this criterion from the analysis to determine the relative importance of the other 4 questions. In this analysis, “Importance” of the scientific question and “Robustness” were most closely linked to overall value (**Supporting Figure S6b**).

Should an article-level correlation of approximately 0.5 between RCR and expert reviewer scores be considered reliable? This level of correspondence is similar to that previously measured between bibliometric indicators and reviewer scores (5,14). Given the partially overlapping datasets used here, we were able to calculate the correlation of expert reviewer scores with one another. The Pearson correlation coefficient between log-transformed F1000 scores and those from the STPI review was 0.35. In addition, the correlation of scores within a survey can be determined with statistical resampling. We selected papers with three reviews from the STPI and NIH IRP surveys. The order of reviewers was randomly shuffled, and the correlation coefficient between the first reviewer’s score and the mean of the other two scores was determined and recorded. This process was repeated 10,000 times for each dataset. The distributions of the 10,000 recorded correlation coefficients are shown in **Supporting Figure S7**. This approach demonstrated an internal correlation of 0.32 for STPI reviews and 0.44 for the NIH IRP reviews. These values are similar to the correlation between RCR and each set of review scores. These internal correlations between reviewer scores likely represent an estimate of the degree to which it is possible for bibliometrics to correspond to expert opinions. Thus, RCR agrees with expert opinion scores as well as experts agree with one another.

### Ranking invariance of RCR

One desirable property in a bibliometric indicator is that of ranking invariance (8,16,17). A citation metric is ranking invariant if, when two groups of articles are being compared using that indicator, their relative ranking does not change if the groups are inflated by the same amount of uncited papers (17). RCR is ranking invariant under two cases: when the two comparison groups are the same size and the same absolute number of uncited papers is added to each group, and when the two comparison groups are different sizes and the same proportion of uncited papers is added to each comparison group.

For the first case (two groups of the same size, termed groups *I* and *J*, where group *I* has the greater RCR), the group RCRs are described by the following inequality:

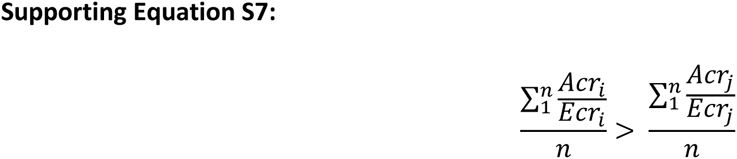

Where *i* and *j* are the individual papers from groups *I* and *J*, and *n* is the number of papers in these groups. Adding *k* > 0 papers to each group, each with a constant RCR of *a* ≥ 0 (equal to 0 for uncited papers) yields the following inequality:

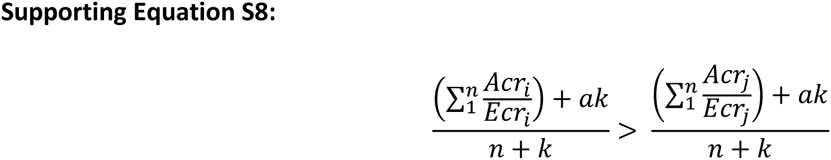

This simplifies to Supporting Equation S9, demonstrating ranking invariance under this condition:

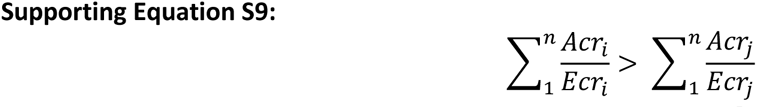

For the second case (two groups of unequal sizes, termed groups *I* and *J*, where group *I* has the greater RCR), the group RCRs are described by the following inequality:

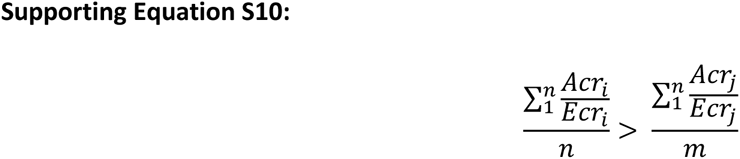

Where *n* is the number of papers in group *I* and *m* is the number of papers in group *J*. Adding the same proportion *k* > 0 of papers to each group, each with a constant RCR of *a* ≥ 0 (equal to 0 for uncited papers) yields the following inequality:

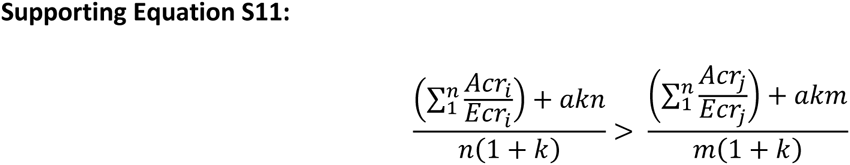

This simplifies back to **Supporting Equation S10**, demonstrating ranking invariance under this condition as well. Note that while uncited papers correspond *a* = 0, any positive RCR *a* could be substituted and ranking invariance would hold.

### Susceptibility to gaming

Since willingness to game Impact Factors is so prevalent (18,19), it stands to reason that some researchers may be tempted to game their RCRs. Self-citation seems to be the most obvious route for boosting the numerator, which is a limitation for citation metrics in general. Is RCR susceptible to gaming of the denominator? Consider this thought experiment: a researcher under career pressure seeks to boost the RCR of one of his papers by lowering its denominator. Is this feasible? In an extreme example, he may publish a new paper citing his previous article, along with 40 others in journals with Impact Factors of 1.0. Effects for a real article published in 2008 with an RCR close to 1.0 are shown in **Supporting Table S5**. The effect of this egregious example is equivalent to only a single additional citation, indicating that even obvious attempts to weigh the cocitation network have, at best, a marginal effect. Obviously, newer articles would be more susceptible to manipulation, but it is unlikely that a researcher would be willing to so overtly attempt to game the metric; much more likely is an attempt to preferentially co-cite related articles in journals at the lower end of normal for the researcher’s field (but not an order of magnitude like our thought experiment). This less obvious form of manipulation would reward the equivalent of much less than one additional citation, and would not have much of an impact.

### Effects of drifting fields over time

Imagine a field whose citation rate drifted significantly over the course of a decade. This field’s intrinsic citation rate went from 6 to 4 over the course of 10 years, and new citations to previously published articles declined by the same amount, in keeping with the field. In the example in **Supporting Table S6**, yearly citations (“Cites”) and their contributions toward FCR (“FCR (Yr.)”) were added as the field’s citation rate declined, contributing lower values to FCR as time went on. The changes in CPY and FCR are shown as separate columns; because these are based on cumulative metrics, their decline lags behind the field as a whole. New article contributions to the co-citation network were modeled with a linear relationship based on empirical results (**Figure 1**). Despite the substantial 33% reduction in the citation rates to this article and its field, the ratio between CPY and FCR is nearly unchanged.

### Investigator-level bibliometrics over long periods

To test the degree to which investigator-level metrics, as shown in **Figure 8** and **Table 2**, are stable over long periods, the analysis was repeated with the 2002-2014 dataset used for the Field Citation Rate stability analysis in **Figure 2**. As in **Figure 8**, investigators with continual R01 funding through the entire window (2002-2013) were selected for inclusion, to rule out additional variance from a catastrophic loss of funding. Rather than comparing two adjacent 4-year windows, two 2-year windows (2002-2003 and 2012-2013) were used spanning a decade. Correlations comparing these two periods are shown in **Supporting Table S7**. Investigator-level correlations are present at this longer time frame, although as expected over a longer time window, at a somewhat lower level. Because this analysis uses two 2-year windows rather than two 4-year windows, this result may underestimate the size of the effect compared to **Figure 8** and **Table 2**.

**Supporting Figure S1.**
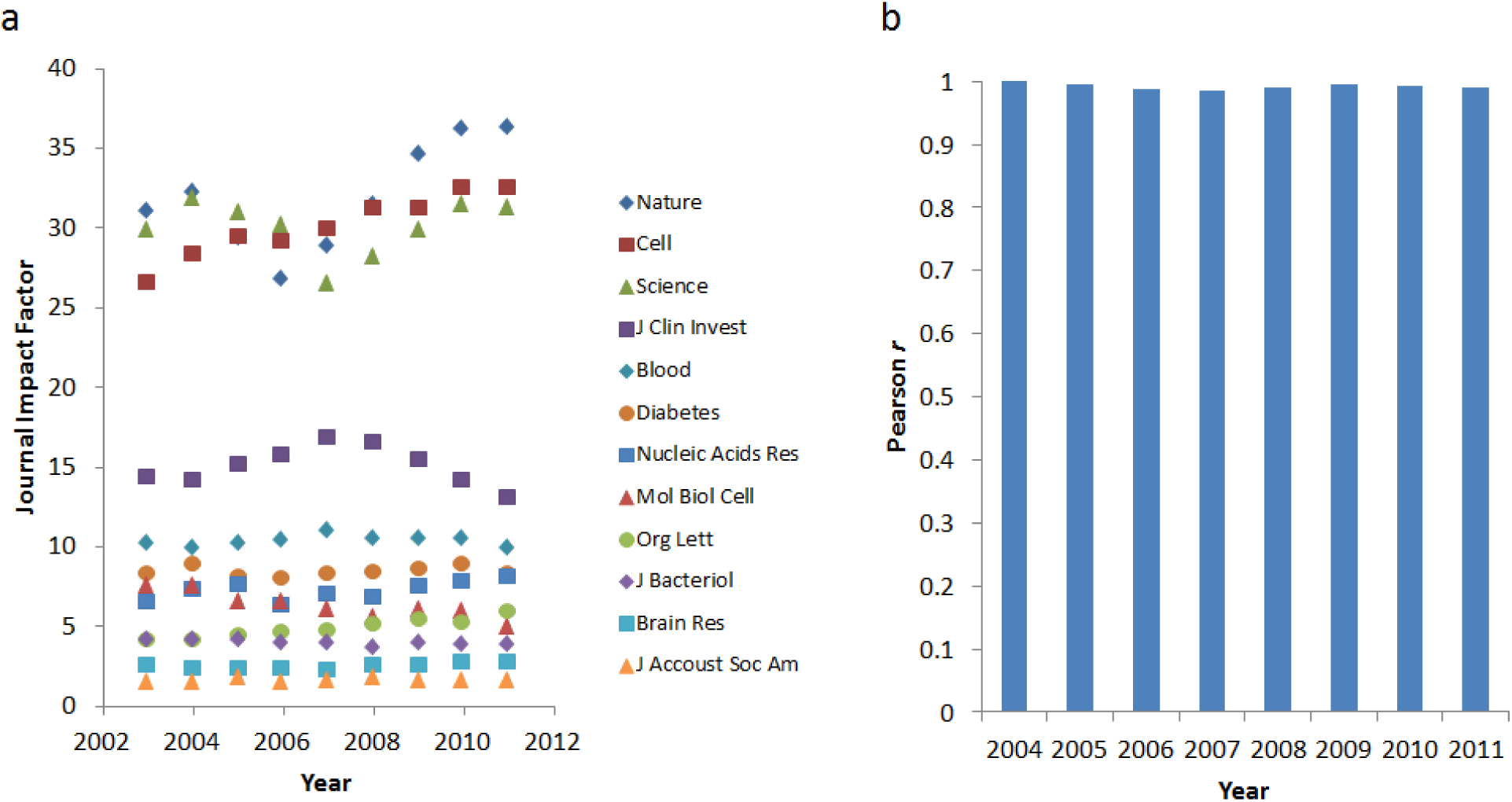
Journal impact factor stability over time. (a) Journal Impact Factors for 12 selected journals from 2003 to 2011. (b) Pearson correlation coefficients ***r*** of the Journal Impact Factors for these 12 journals in 2003 vs. each of their respective Impact Factors in subsequent years. In each case, ***r*** is over 0.9.

**Supporting Figure S2.**
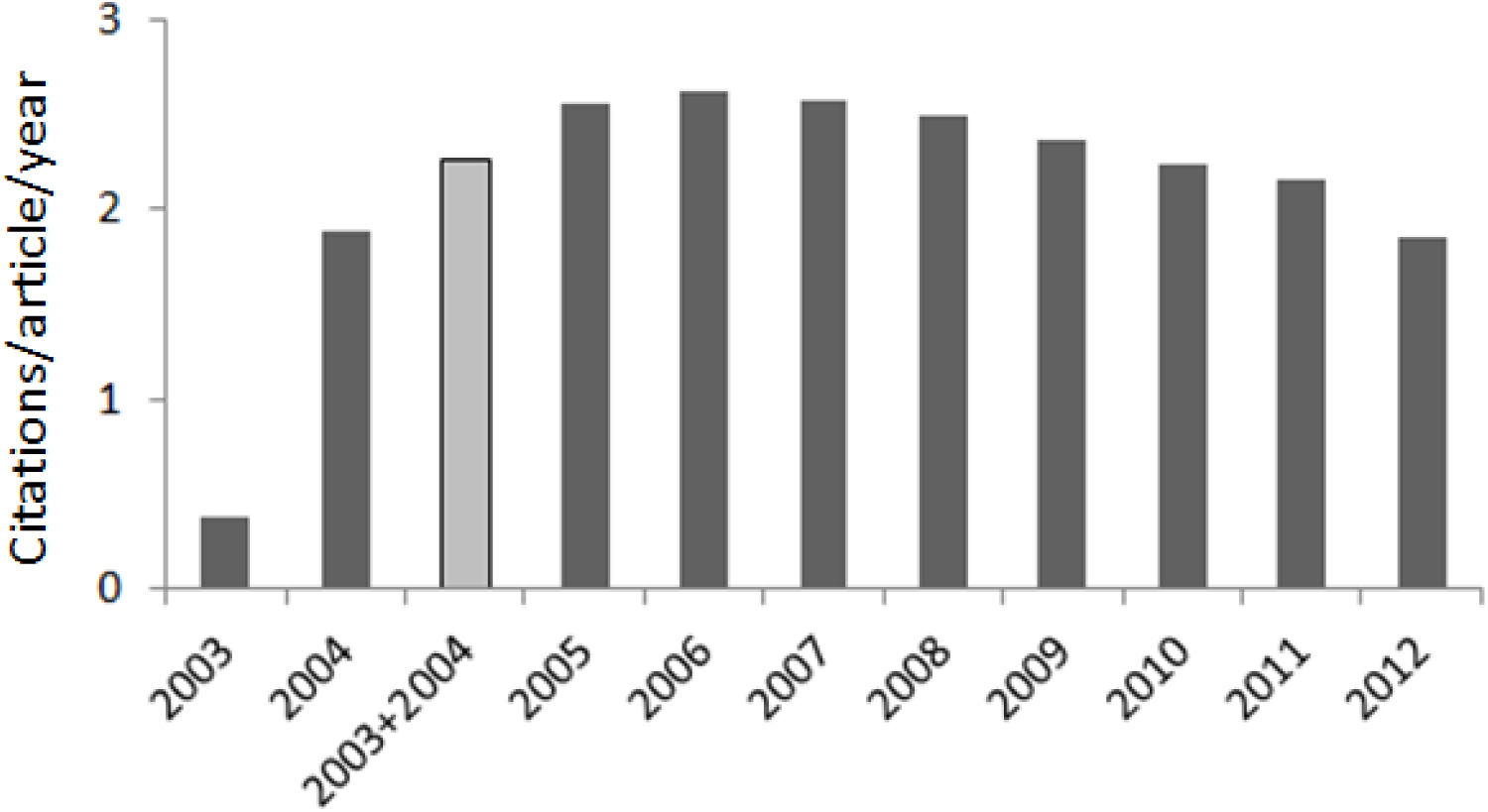
Mean citations accrued each year for 608,058 papers published in 2003 appearing in the same journals as NIH-funded publications. Adding the values for 2003 and 2004 gives a value (2.26 citations per publication per year) close to the mean citations per year of the following years (2.36). Although these values may seem low, they are both similar to the global 2013 Aggregate Impact Factor metric for journals appearing in the Biology subcategory (2.56), which is also measured in citations per paper per year.

**Supporting Figure S3.**
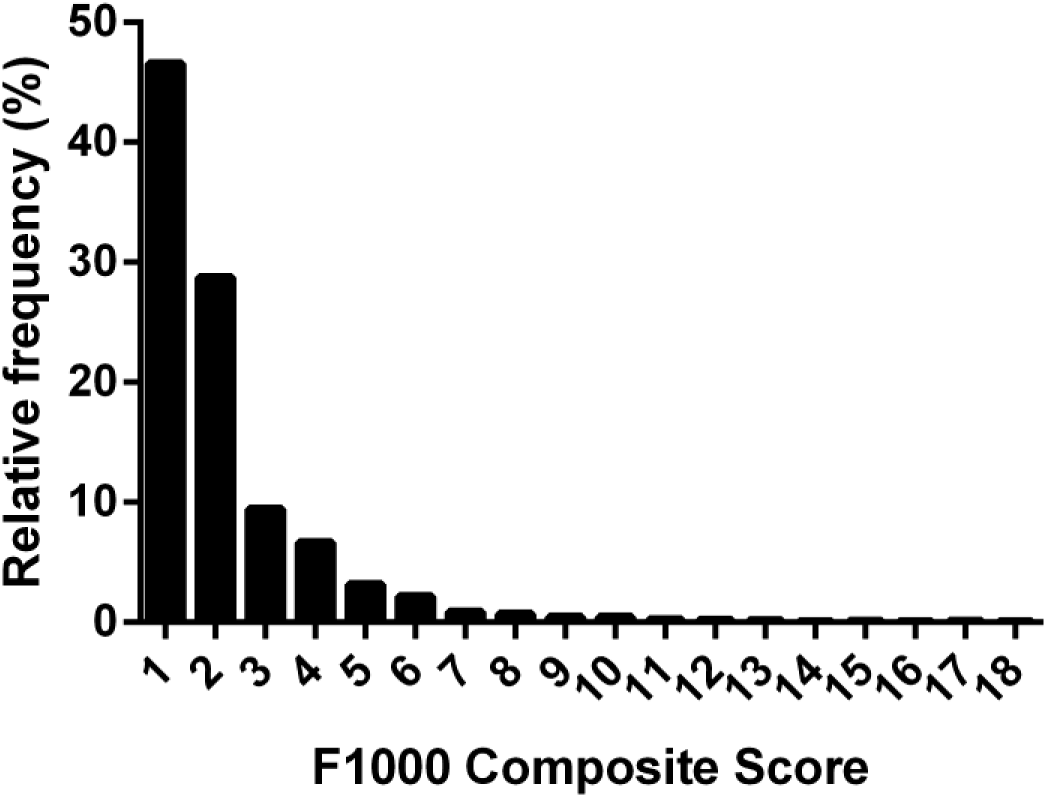
Distribution of Faculty of 1000 scores for 2193 R01-funded papers from 2009.

**Supporting Figure S4.**
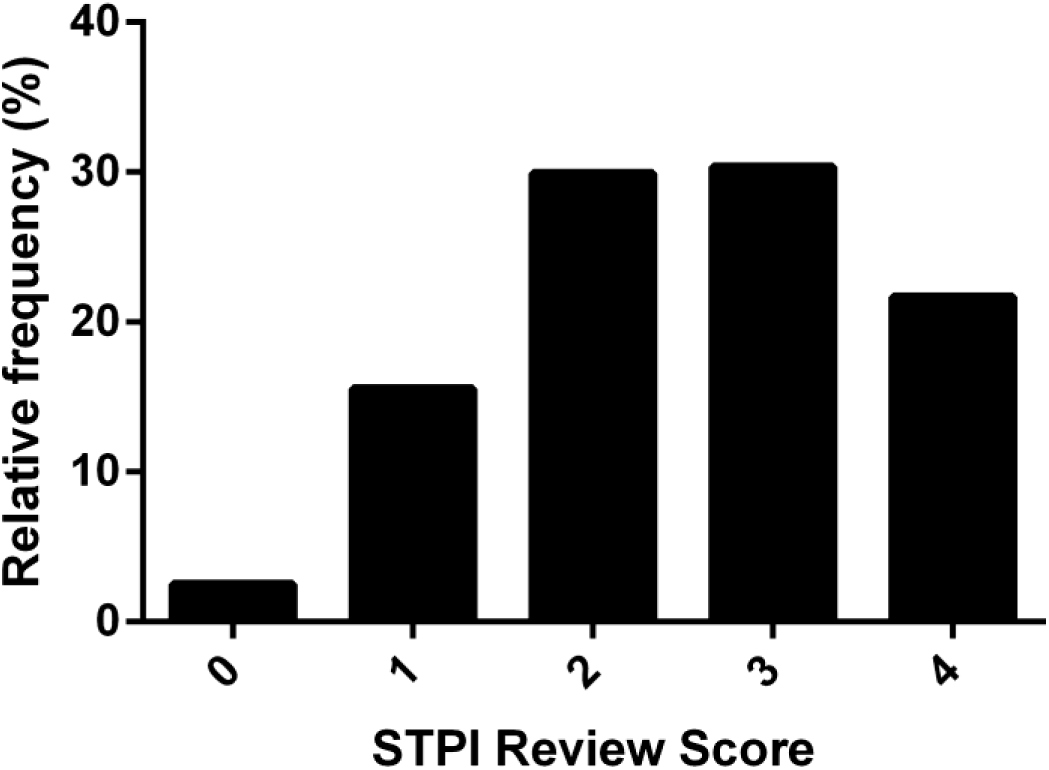
Distribution of scores from the Science and Technology Policy Institute survey.

**Supporting Figure S5.**
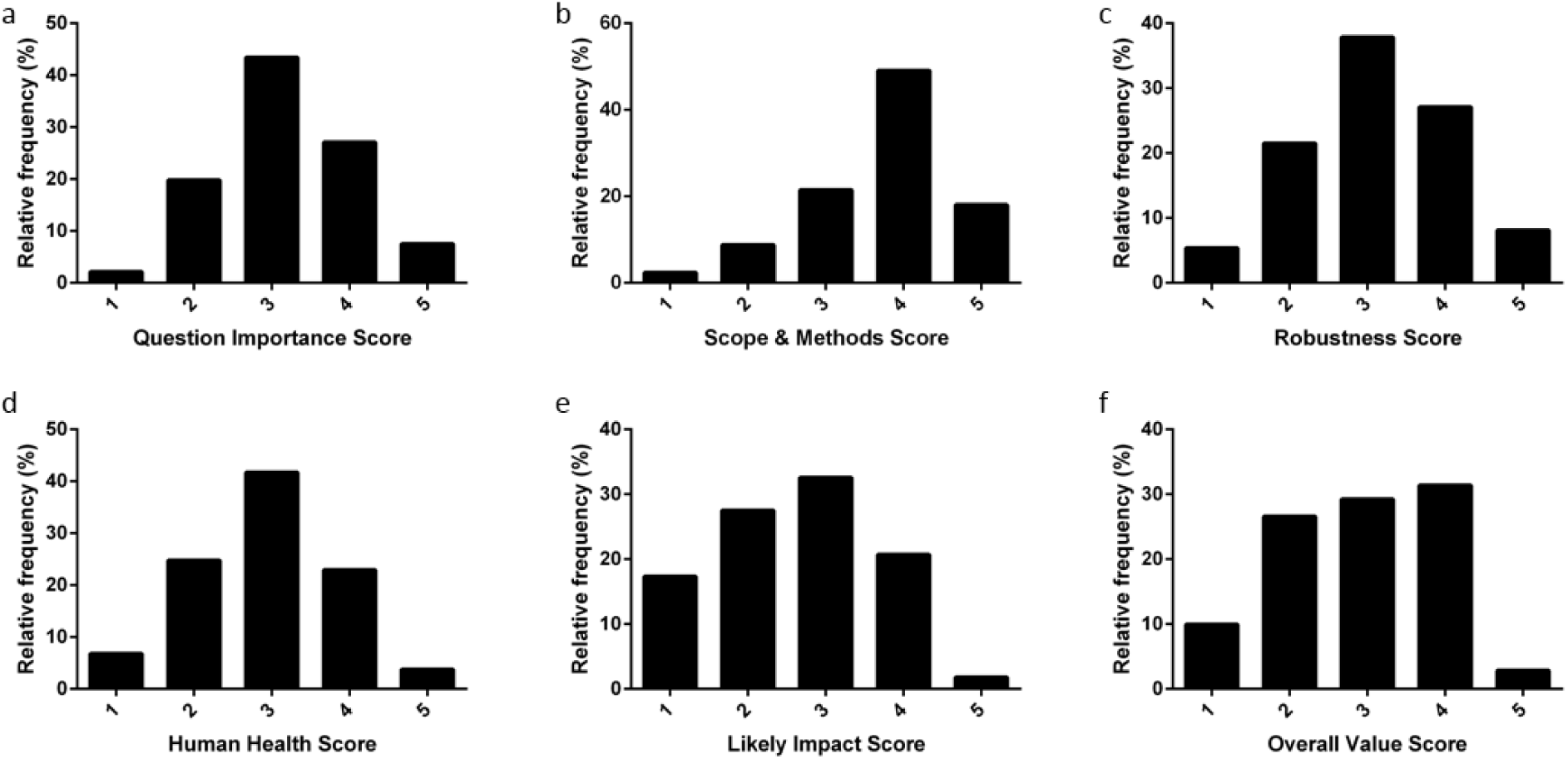
Summary of NIH IRP reviewer responses to post-publication peer review questions. Distribution of ratings to the questions: (a) Rate whether the question being addressed is important to answer. (b) Rate whether you agree that the methods are appropriate and the scope of the experiments adequate. (c) Rate how robust the study is based on the strength of the evidence presented. (d) Rate the likelihood that the results could ultimately have a substantial positive impact on human health outcomes. (e) Rate the impact that the research is likely to have or has already had. (f) Provide your overall evaluation of the value and impact of this publication.

**Supporting Figure S6.**
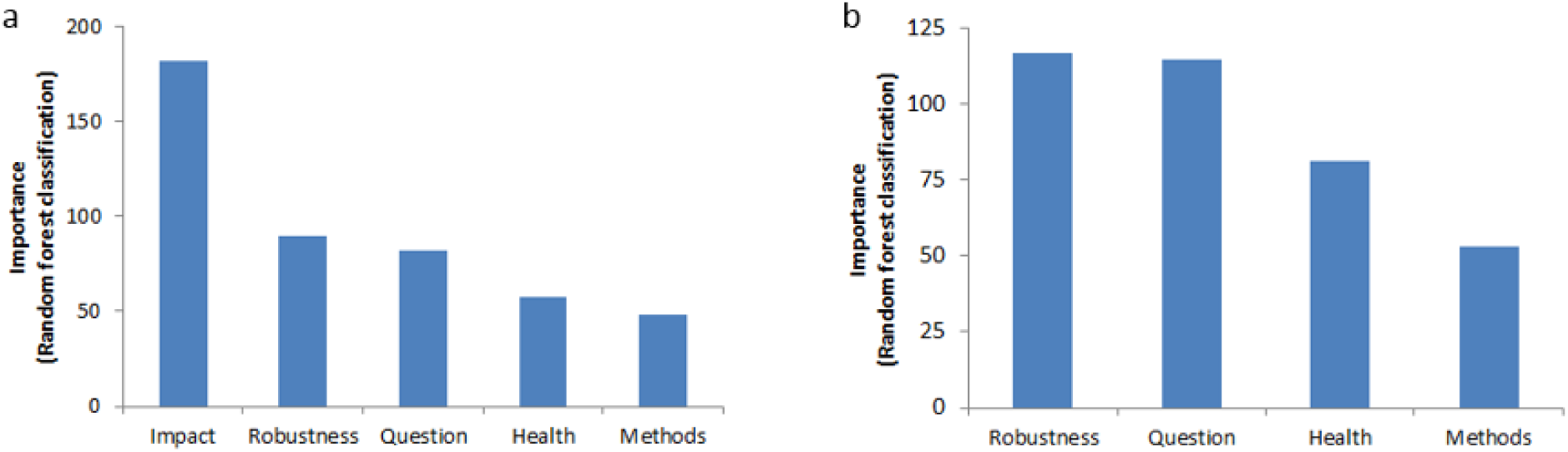
Inferred criteria used by NIH IRP reviewers to rate overall value and impact of a publication. (a) Criteria most strongly linked to assessments of “overall value”, measured with Random Forest classification. Values indicate the mean decrease in Gini coefficient. (b) Criteria most strongly linked to assessments of “overall value”, excluding “likely impact”, and measured with Random Forest classification.

**Supporting Figure S7.**
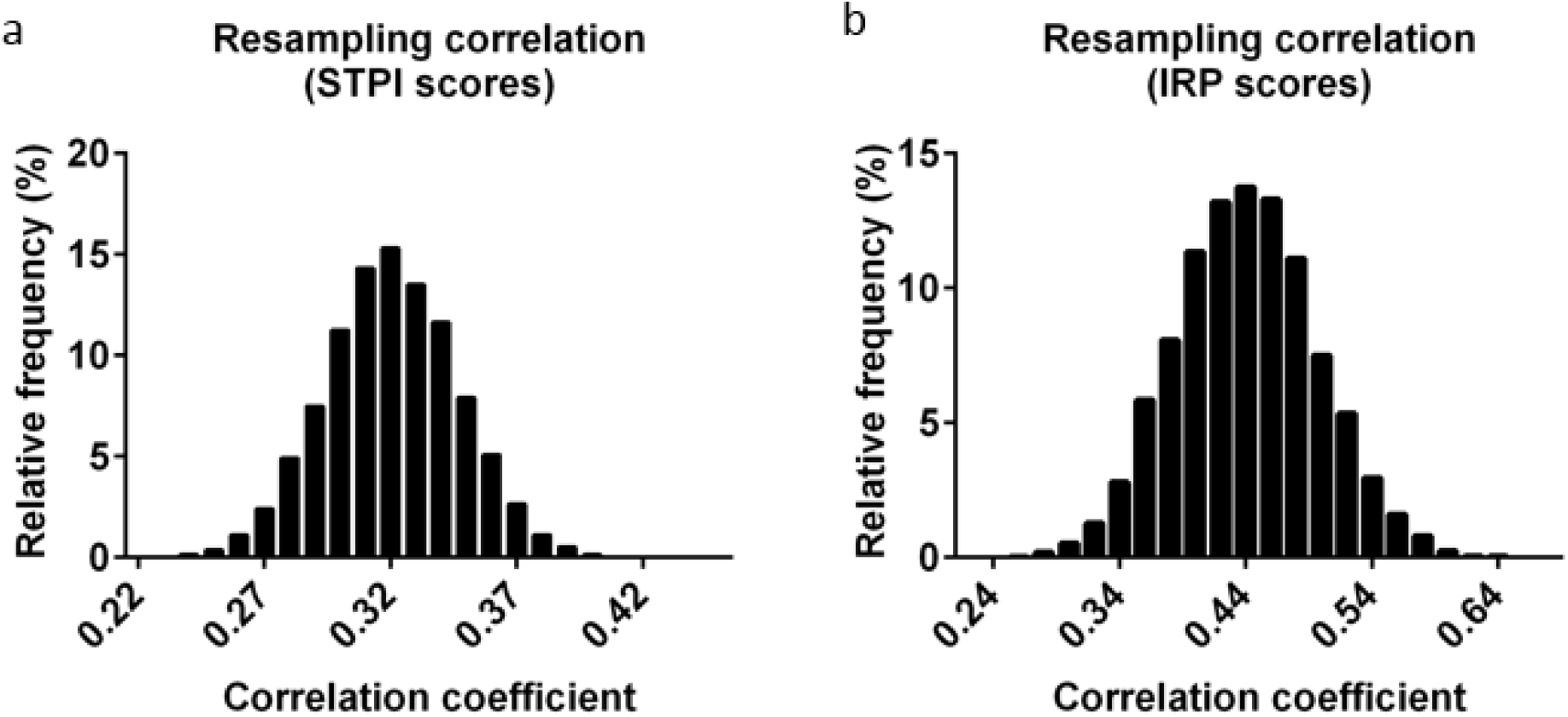
Internal correlation of post-publication peer review scores is similar to correlation between RCR and review scores. (a) Pearson correlation coefficients (***r***) of one randomly chosen reviewer score vs. the mean of the other two scores for that paper, determined by statistical resampling. Distribution of correlation coefficients determined by resampling from the STPI dataset (10,000 repetitions, mean ***r*** = 0.32). (b) Pearson correlation coefficients (***r***) of one randomly chosen reviewer score vs. the mean of the other two scores for that paper, determined by statistical resampling. Distribution of correlation coefficients determined by resampling from the IRP dataset (10,000 repetitions, mean ***r*** = 0.44).

**Supporting Figure S8.**
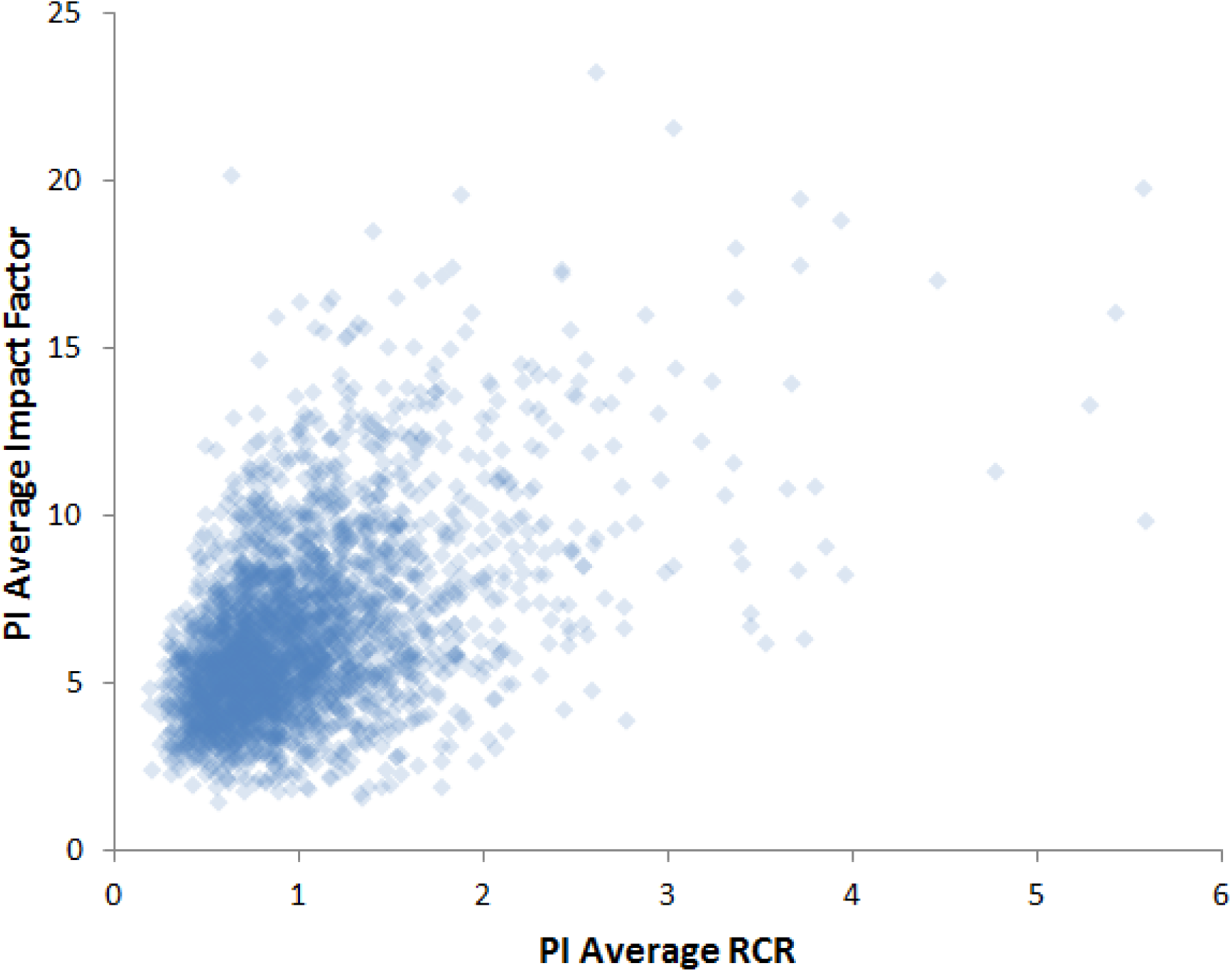
Correlation of the average of different investigators’ article RCRs vs. the average of the Journal Impact Factors in which they published. Some investigators published very influential articles (high RCR) in lower-profile venues (low JIF) and vice versa. R^2^ = 0.23

**Supporting Table S1.**
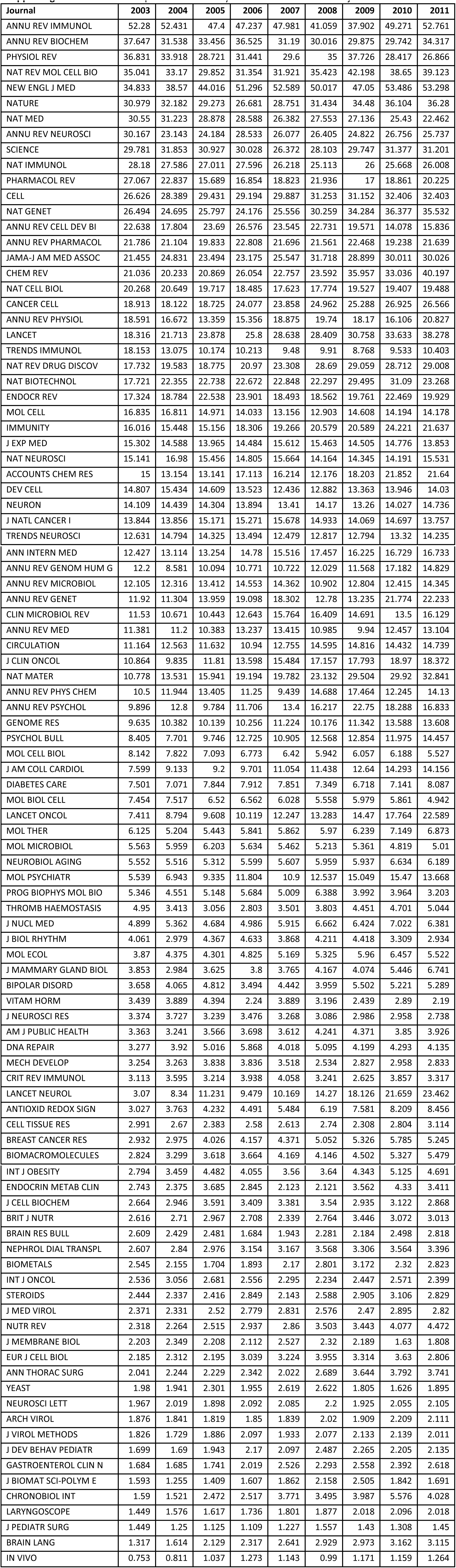
Journal Impact Factor stability over time for 100 selected journals.

**Supporting Table S2.** Regression coefficients (ACR on FCR, through 2012), for ordinary least squares linear regression (OLS) or quantile regression to the median (QR) for papers with concurrent R01 funding.

| Year | Slope (OLS) | Intercept (OLS) | Slope (QR) | Intercept (QR) |
| --- | --- | --- | --- | --- |
| 2003 | 0.497 | 1.947 | 0.355 | 0.870 |
| 2004 | 0.531 | 1.877 | 0.344 | 1.090 |
| 2005 | 0.593 | 1.645 | 0.384 | 0.914 |
| 2006 | 0.608 | 1.471 | 0.385 | 0.967 |
| 2007 | 0.643 | 1.411 | 0.381 | 1.091 |
| 2008 | 0.772 | 0.711 | 0.446 | 0.681 |
| 2009 | 0.768 | 0.704 | 0.469 | 0.542 |
| 2010 | 0.739 | 0.657 | 0.490 | 0.156 |
| 2011 | 0.780 | 0.120 | 0.498 | 0.000 |

One of the criticisms about Impact Factor is that it uses the mean, rather than median, of a skewed distribution (20); for RCR, quantile regression can be used to benchmark articles to the median citation performance in the benchmark group, rather than the mean.

**Supporting Table S3.** Correlation coefficients (*r*) between the log-transformed RCR values generated with the <u>three different methods for estimating field citation rates</u> with co-citation networks.

| Methods compared | Pearson $r$ |
| --- | --- |
| Cited ~ Citing | 0.928529 |
| Cited ~ Co-cited | 0.95274 |
| Citing ~ Co-Cited | 0.959203 |

**Supporting Table S4.** Correlation between ACR vs. ECR.

| Method used for expected citation rate | Pearson $r$ |
| --- | --- |
| Cited | 0.292319 |
| Citing | 0.250813 |
| Co-cited | 0.180248 |

**Supporting Table S5.** Effects of an attempt to game the denominator by introducing low Impact-Factor articles (40 articles of Impact Factor 1.0) to the co-citation network to a real article close to the average RCR.

|  | Pre-gaming | Post-gaming |
| --- | --- | --- |
| Co-citation network size | 1238 | 1278 |
| FCR | 10.2 | 9.9 |
| ECR | 7.9 | 7.7 |
| CPY | 7.6 | 7.8 |
| RCR change from +1 cite | 0.962 | 0.987 |
| RCR change from ΔFCR | 0.962 | 0.987 |
| RCR total change | 0.962 | 1.01 |

**Supporting Table S6.** Effects on CPY:FCR ratio of a field with a citation rate that drifts by a factor of 1.5 over the course of 10 years.

| Year | Cites | CPY | FCR (Yr.) | FCR | CPY : FCR |
| --- | --- | --- | --- | --- | --- |
| 2003 | 0 | - | 6 | - | - |
| 2004 | 6 | 6 | 6 | 6 | 1.00 |
| 2005 | 6 | 6 | 6 | 6 | 1.00 |
| 2006 | 6 | 6 | 6 | 6 | 1.00 |
| 2007 | 5 | 5.75 | 5 | 5.782609 | 0.99 |
| 2008 | 5 | 5.6 | 5 | 5.642857 | 0.99 |
| 2009 | 5 | 5.5 | 5 | 5.545455 | 0.99 |
| 2010 | 4 | 5.285714 | 4 | 5.378378 | 0.98 |
| 2011 | 4 | 5.125 | 4 | 5.243902 | 0.98 |
| 2012 | 4 | 5 | 4 | 5.133333 | 0.97 |

**Supporting Table S7.** Summary of investigator-level bibliometric measures and their stability from two 2-year periods spanning a decade (2002-2003 and 2012-2013, PIs with more than 5 articles in each period).

| Measure | Pearson *r* (of log-transformed values) |
| --- | --- |
| Field Citation Rate | 0.83 |
| Journal Citation Rate | 0.64 |
| Article Citation Rate | 0.48 |
| RCR | 0.40 |

## References

1. Couzin-Frankel J. Shaking up science. Science [Internet]. 2013 Jan 25;339(6118):386–9.

2. Hirsch JE. An index to quantify an individual’s scientific research output. Proc Natl Acad Sci U S A [Internet]. 2005 Nov 15;102(46):16569–72.

3. Seglen PO. Why the impact factor of journals should not be used for evaluating research. BMJ [Internet]. 1997 Feb 15;314(7079):497–497.

4. Not-so-deep impact. Nature [Internet]. Nature Publishing Group; 2005 Jun 23;435(7045):1003–4.

5. The maze of impact metrics. Nature [Internet]. 2013 Oct 16;502(7471):271–271.

6. Pulverer B. Impact fact-or fiction? EMBO J [Internet]. EMBO Press; 2013 Jun 12;32(12):1651–2.

7. Stallings J, Vance E, Yang J, Vannier MW, Liang J, Pang L, et al. Determining scientific impact using a collaboration index. Proc Natl Acad Sci U S A [Internet]. 2013 Jun 11;110(24):9680–5.

8. Garfield E. The history and meaning of the journal impact factor. JAMA [Internet]. American Medical Association; 2006 Jan 4;295(1):90–3.

9. Stringer MJ, Sales-Pardo M, Nunes Amaral LA. Effectiveness of journal ranking schemes as a tool for locating information. PLoS One [Internet]. Public Library of Science; 2008 Jan 27;3(2):e1683.

10. Misteli T. Eliminating the impact of the Impact Factor. J Cell Biol [Internet]. 2013 May 27;201(5):651–2.

11. Johnston M. We have met the enemy, and it is us. Genetics [Internet]. 2013 Aug 1;194(4):791–2.

12. Moed HF, Burger WJM, Frankfort JG, Van Raan AFJ. The use of bibliometric data for the measurement of university research performance. Res Policy [Internet]. 1985 Jun;14(3):131–49.

13. Zitt M, Small H. Modifying the journal impact factor by fractional citation weighting: The audience factor. J Am Soc Inf Sci Technol [Internet]. 2008 Sep;59(11):1856–60.

14. Bornmann L, Leydesdorff L. The validation of (advanced) bibliometric indicators through peer assessments: A comparative study using data from InCites and F1000. J Informetr [Internet]. 2013 Apr;7(2):286–91.

15. Opthof T, Leydesdorff L. Caveats for the journal and field normalizations in the CWTS (“Leiden”) evaluations of research performance. J Informetr [Internet]. 2010 Jul;4(3):423–30.

16. van Raan AFJ, van Leeuwen TN, Visser MS, van Eck NJ, Waltman L. Rivals for the crown: Reply to Opthof and Leydesdorff. J Informetr [Internet]. 2010 Jul;4(3):431–5.

17. Waltman L, van Eck NJ, van Leeuwen TN, Visser MS, van Raan AFJ. Towards a new crown indicator: Some theoretical considerations. J Informetr [Internet]. 2011 Jan;5(1):37–47.

18. Waltman L, Yan E, van Eck NJ. A recursive field-normalized bibliometric performance indicator: an application to the field of library and information science. Scientometrics [Internet]. Akadémiai Kiadó, copublished with Springer Science*Business Media B.V., Formerly Kluwer Academic Publishers B.V.; 2011 Jul 10

19. Bornmann L, Marx W. How to evaluate individual researchers working in the natural and life sciences meaningfully? A proposal of methods based on percentiles of citations. Scientometrics [Internet]. 2013 Oct 23;98(1):487–509.

20. Bergstrom CT, West JD, Wiseman MA. The Eigenfactor metrics. J Neurosci [Internet]. 2008 Nov 5;28(45):11433–4.

21. Bergstrom CT, West JD. Assessing citations with the Eigenfactor metrics. Neurology [Internet]. 2008 Dec 2;71(23):1850–1.

22. Moed HF. Measuring contextual citation impact of scientific journals. J Informetr [Internet]. 2010 Jul;4(3):265–77.

23. Bollen J, Van de Sompel H, Hagberg A, Chute R. A principal component analysis of 39 scientific impact measures. PLoS One [Internet]. Public Library of Science; 2009 Jan 29;4(6):e6022.

24. Waltman L, van Eck NJ, van Leeuwen TN, Visser MS. Some modifications to the SNIP journal impact indicator. J Informetr [Internet]. 2013 Apr;7(2):272–85.

25. Waltman L, van Eck NJ. A systematic empirical comparison of different approaches for normalizing citation impact indicators. J Informetr [Internet]. 2013 Oct;7(4):833–49.

26. Wang D, Song C, Barabàsi A-L. Quantifying long-term scientific impact. Science [Internet]. 2013 Oct 4;342(6154):127–32.

27. Radicchi F, Fortunato S, Castellano C. Universality of citation distributions: toward an objective measure of scientific impact. Proc Natl Acad Sci U S A [Internet]. 2008 Nov 11;105(45):17268–72.

28. Stringer MJ, Sales-Pardo M, Nunes Amaral LA. Statistical validation of a global model for the distribution of the ultimate number of citations accrued by papers published in a scientific journal. J Am Soc Inf Sci Technol JASIST [Internet]. 2010 Jul;61(7):1377–85.

29. Walker D, Xie H, Yan K-K, Maslov S. Ranking scientific publications using a model of network traffic. J Stat Mech Theory Exp [Internet]. 2007 Jun 14;2007(06):P06010–P06010.

30. Crous CJ. Judge research impact on a local scale. Nature [Internet]. 2014 Sep 4;513(7516):7.

31. Radicchi F, Castellano C. Testing the fairness of citation indicators for comparison across scientific domains: The case of fractional citation counts. J Informetr [Internet]. 2012 Jan;6(1):121–30.

32. Jeong H, Néda Z, Barabàsi AL. Measuring preferential attachment in evolving networks. Europhys Lett [Internet]. 2003 Feb 2;61(4):567–72.

33. Eom Y-H, Fortunato S. Characterizing and modeling citation dynamics. PLoS One [Internet]. Public Library of Science; 2011 Jan 22;6(9):e24926.

34. Small H. Co-citation in the scientific literature: A new measure of the relationship between two documents. J Am Soc Inf Sci [Internet]. 1973 Jul;24(4):265–9.

35. Mir R, Karim S, Kamal MA, Wilson CM, Mirza Z. Conotoxins: Structure, Therapeutic Potential and Pharmacological Applications. Curr Pharm Des [Internet]. 2015 Nov 24;

36. Lewis J, Ossowski S, Hicks J, Errami M, Garner HR. Text similarity: an alternative way to search MEDLINE. Bioinformatics [Internet]. 2006 Sep 15;22(18):2298–304.

37. Breiman L. Random Forests. Mach Learn [Internet]. Kluwer Academic Publishers;;45(1):5–32.

38. Rousseau R, Leydesdorff L. Simple arithmetic versus intuitive understanding:The case of the impact factor [Internet]. ISSI Newsletter. ISSI; 2011. p. 10–4.

39. Glanzel W, Moed HF. Opinion paper: thoughts and facts on bibliometric indicators. Scientometrics [Internet]. 2012 Nov 21;96(1):381–94.

40. Bornmann L, Haunschild R. Relative Citation Ratio (RCR): A first empirical attempt to study a new field-normalized bibliometric indicator. 2015 Nov 25

41. Ke Q, Ferrara E, Radicchi F, Flammini A. Defining and identifying Sleeping Beauties in science. Proc Natl Acad Sci U S A [Internet]. 2015 Jun 16;112(24):7426–31.

42. Hicks D, Wouters P, Waltman L, de Rijcke S, Rafols I. Bibliometrics: The Leiden Manifesto for research metrics. Nature [Internet]. 2015 Apr 23;520(7548):429–31.

43. Ribas S, Ueda A, Santos RLT, Ribeiro-Neto B, Ziviani N. Simplified Relative Citation Ratio for Static Paper Ranking: UFMG/LATIN at WSDM Cup 2016. 2016 Mar 3

44. Page L, Brin S, Motwani R, Winograd T. The PageRank Citation Ranking: Bringing Order to the Web. [Internet]. Stanford InfoLab; 1999.

45. Chen P, Xie H, Maslov S, Redner S. Finding scientific gems with Google’s PageRank algorithm. J Informetr [Internet]. 2007 Jan;1(1):8–15.

46. Yan E, Ding Y. Discovering author impact: A PageRank perspective. Inf Process Manag [Internet]. 2011 Jan;47(1):125–34.

47. Leydesdorff L, Bornmann L. The operationalization of “fields” as WoS subject categories (WCs) in evaluative bibliometrics: The cases of “library and information science” and “science & technology studies.” J Assoc Inf Sci Technol [Internet]. 2015 Feb 8

48. Price DDS. A general theory of bibliometric and other cumulative advantage processes. J Am Soc Inf Sci [Internet]. 1976;27(5-6):292–306.

49. Sullivan MX. Synthetic Culture Media and the Biochemistry of bacterial Pigments. J Med Res [Internet]. 1905 Nov;14(1):109–60.

50. Institute of Medicine NA of S and NA of E. Facilitating Interdisciplinary Research [Internet]. Washington, D.C.: National Academies Press; 2004.

51. Huerta MF, Farber GK, Wilder EL, Kleinman D V, Grady PA, Schwartz DA, et al. NIH Roadmap interdisciplinary research initiatives. PLoS Comput Biol [Internet]. Public Library of Science; 2005 Nov 25;1(6):e59.

52. Mansilla R, Koppen E, Cocho G, Miramontes P. On the behavior of journal impact factor rank-order distribution. J Informetr [Internet]. 2007 Apr;1(2):155–60.

53. Egghe L. Mathematical derivation of the impact factor distribution. J Informetr [Internet]. 2009 Oct;3(4):290–5.

54. Casadevall A, Fang FC. Causes for the persistence of impact factor mania. MBio [Internet]. 2014 Jan;5(2):e00064–14.

55. Shaw D. The prisoners’ dilemmas: Authorship guidelines and impact factors: between a rock and a hard place. EMBO Rep [Internet]. 2014 Jun;15(6):635–7.

56. Bertuzzi S, Drubin DG. No shortcuts for research assessment. Mol Biol Cell [Internet]. 2013 May 15;24(10):1505–6.

57. Suhrbier A, Poland GA. Are Impact Factors corrupting truth and utility in biomedical research? Vaccine [Internet]. 2013 Dec 9;31(51):6041–2.

58. Alberts B. Impact factor distortions. Science [Internet]. 2013 May 17;340(6134):787.

59. Schekman R, Patterson M. Reforming research assessment. Elife [Internet]. eLife Sciences Publications Limited; 2013 Jan 16;2:e00855.

60. Vinkler P. Relations of relative scientometric indicators. Scientometrics [Internet]. 2003;58(3):687–94.

61. Rand DG, Pfeiffer T. Systematic differences in impact across publication tracks at PNAS. PLoS One [Internet]. Public Library of Science; 2009 Jan 1;4(12):e8092.

62. Wilhite AW, Fong EA. Scientific publications. Coercive citation in academic publishing. Science [Internet]. American Association for the Advancement of Science; 2012 Feb 3;335(6068):542–3.

63. Krell F-T. Should editors influence journal impact factors? Learn Publ [Internet]. 2010 Jan 1;23(1):59–62.

64. Salganik MJ, Dodds PS, Watts DJ. Experimental study of inequality and unpredictability in an artificial cultural market. Science [Internet]. American Association for the Advancement of Science; 2006 Feb 10;311(5762):854–6.

65. Liao H, Xiao R, Cimini G, Medo M. Network-driven reputation in online scientific communities. PLoS One [Internet]. Public Library of Science; 2014 Jan 2;9(12):e112022.

66. Danthi N, Wu CO, Shi P, Lauer M. Percentile ranking and citation impact of a large cohort of National Heart, Lung, and Blood Institute-funded cardiovascular R01 grants. Circ Res [Internet]. 2014 Feb 14;114(4):600–6.

67. Kaltman JR, Evans FJ, Danthi NS, Wu CO, DiMichele DM, Lauer MS. Prior publication productivity, grant percentile ranking, and topic-normalized citation impact of NHLBI cardiovascular R01 grants. Circ Res [Internet]. 2014 Sep 12;115(7):617–24.

68. Mervis J. Peering into peer review. Science [Internet]. 2014 Feb 7;343(6171):596–8.

69. DePellegrin TA, Johnston M. An Arbitrary Line in the Sand: Rising Scientists Confront the Impact Factor. Genetics [Internet]. 2015 Nov 12;201(3):811–3.

70. Contopoulos-Ioannidis DG, Alexiou GA, Gouvias TC, Ioannidis JPA. Medicine. Life cycle of translational research for medical interventions. Science [Internet]. 2008 Sep 5;321(5894):1298–9.

71. Weber GM. Identifying translational science within the triangle of biomedicine. J Transl Med [Internet]. 2013 Jan;11(1):126.

72. Garfield E. Citation analysis as a tool in journal evaluation. Science [Internet]. 1972 Nov 3;178(4060):471–9.

73. Hutchins BI, Yuan X, Anderson JM, Santangelo GM. Relative Citation Ratio (RCR): A new metric that uses citation rates to measure influence at the article level [Internet]. bioRxiv. Cold Spring Harbor Labs Journals; 2015 Oct.

74. Rose S, Engel D, Cramer N, Cowley W. Text Mining: Autometic keyword extraction from individual documents. Berry MW, Kogan J, editors. Chichester, UK: John Wiley & Sons, Ltd; 2010.

75. Feinerer I, Hornik K, Meyer D. Text Mining Infrastructure in R. J Stat Softw [Internet]. 2008 Mar 31;25(5):1–54.

## Supporting References

1. Schubert A, Glänzel W, Braun T. Relative indicators of publication output and citation impact of european physics research, 1978–1980. Czechoslov J Phys [Internet]. 1986 Jan;36(1):126–9.

2. Vinkler P. Relations of relative scientometric indicators. Scientometrics [Internet]. 2003;58(3):687–94.

3. Moed HF, Burger WJM, Frankfort JG, Van Raan AFJ. The use of bibliometric data for the measurement of university research performance. Res Policy [Internet]. 1985 Jun;14(3):131–49.

4. Zitt M, Small H. Modifying the journal impact factor by fractional citation weighting: The audience factor. J Am Soc Inf Sci Technol [Internet]. 2008 Sep;59(11):1856–60.

5. Bornmann L, Leydesdorff L. The validation of (advanced) bibliometric indicators through peer assessments: A comparative study using data from InCites and F1000. J Informetr [Internet]. 2013 Apr;7(2):286–91.

6. Opthof T, Leydesdorff L. Caveats for the journal and field normalizations in the CWTS (“Leiden”) evaluations of research performance. J Informetr [Internet]. 2010 Jul;4(3):423–30.

7. van Raan AFJ, van Leeuwen TN, Visser MS, van Eck NJ, Waltman L. Rivals for the crown: Reply to Opthof and Leydesdorff. J Informetr [Internet]. 2010 Jul;4(3):431–5.

8. Waltman L, van Eck NJ, van Leeuwen TN, Visser MS, van Raan AFJ. Towards a new crown indicator: Some theoretical considerations. J Informetr [Internet]. 2011 Jan;5(1):37–47.

9. Lundberg J. Lifting the crown—citation z-score. J Informetr [Internet]. 2007 Apr;1(2):145–54.

10. Leydesdorff L, Bornmann L. The operationalization of “fields” as WoS subject categories (WCs) in evaluative bibliometrics: The cases of “library and information science” and “science & technology studies.” J Assoc Inf Sci Technol [Internet]. 2015 Feb 8

11. Talley EM, Newman D, Mimno D, Herr BW, Wallach HM, Burns GAPC, et al. Database of NIH grants using machine-learned categories and graphical clustering. Nat Methods [Internet]. Nature Publishing Group, a division of Macmillan Publishers Limited. All Rights Reserved.; 2011 Jun;8(6):443–4.

12. Small H. Co-citation in the scientific literature: A new measure of the relationship between two documents. J Am Soc Inf Sci [Internet]. 1973 Jul;24(4):265–9.

13. Koenker R. Quantile Regression [Internet]. Cambridge University Press; 2005. 349

14. Li X, Thelwall M. F1000, Mendeley and Traditional Bibliometric Indicators. Proc 17th Int Conf Sci Technol Indic Montréal Sci OST [Internet]. 2012;3:541–51.

15. Lal B, Wilson AG, Jonas S, Lee EC, Richards AM, Peña VI. An Outcome Evaluation of the National Institutes of Health (NIH) Director’s Pioneer Award (NDPA) Program, FY 2004-2006. 2012.

16. Glanzel W, Moed HF. Opinion paper: thoughts and facts on bibliometric indicators. Scientometrics [Internet]. 2012 Nov 21;96(1):381–94.

17. Rousseau R, Leydesdorff L. Simple arithmetic versus intuitive understanding:The case of the impact factor [Internet]. ISSI Newsletter. ISSI; 2011. p. 10–4.

18. Wilhite AW, Fong EA. Scientific publications. Coercive citation in academic publishing. Science [Internet]. American Association for the Advancement of Science; 2012 Feb 3;335(6068):542–3.

19. Krell F-T. Should editors influence journal impact factors? Learn Publ [Internet]. 2010 Jan 1;23(1):59–62.

20. Rossner M, van Epps H, Hill E. Show me the data. J Cell Biol [Internet]. 2007 Dec 17;179(6):1091–2.

